# Tetravalent SARS-CoV-2 Neutralizing Antibodies Show Enhanced Potency and Resistance to Escape Mutations

**DOI:** 10.1101/2020.10.31.362848

**Authors:** Shane Miersch, Zhijie Li, Reza Saberianfar, Mart Ustav, James Brett Case, Levi Blazer, Chao Chen, Wei Ye, Alevtina Pavlenco, Maryna Gorelik, Julia Garcia Perez, Suryasree Subramania, Serena Singh, Lynda Ploder, Safder Ganaie, Rita E. Chen, Daisy W. Leung, Pier Paolo Pandolfi, Giuseppe Novelli, Giulia Matusali, Francesca Colavita, Maria R. Capobianchi, Suresh Jain, J.B. Gupta, Gaya K. Amarasinghe, Michael S. Diamond, James Rini, Sachdev S. Sidhu

**Author notes:** The authors contributed equally (Shane Miersch, Zhijie Li).

## Abstract

Neutralizing antibodies (nAbs) hold promise as effective therapeutics against COVID-19. Here, we describe protein engineering and modular design principles that have led to the development of synthetic bivalent and tetravalent nAbs against SARS-CoV-2. The best nAb targets the host receptor binding site of the viral S-protein and its tetravalent versions can block entry with a potency that exceeds the bivalent nAbs by an order of magnitude. Structural studies show that both the bivalent and tetravalent nAbs can make multivalent interactions with a single S-protein trimer, observations consistent with the avidity and potency of these molecules. Significantly, we show that the tetravalent nAbs show much increased tolerance to potential virus escape mutants. Bivalent and tetravalent nAbs can be produced at large-scale and are as stable and specific as approved antibody drugs. Our results provide a general framework for developing potent antiviral therapies against COVID-19 and related viral threats, and our strategy can be readily applied to any antibody drug currently in development.

## INTRODUCTION

As of December 10, 2020, the ongoing COVID-19 viral pandemic has tallied more than 68,000,000 confirmed cases and caused over 1,550,000 deaths (www.who.int). Moreover, the highly infectious nature of the disease has imposed severe global economic hardship due to the need for social distancing and lockdown measures. A number of repurposed drugs have shown only limited or uncertain efficacy against COVID-19 (Beigel et al., 2020; Boulware et al., 2020). Although licensed vaccines are now emerging (Poland et al., 2020), their effectiveness across demographics, their availability worldwide, and how well they will be adopted by the public at large remains an unknown. Almost certainly, COVID-19 will remain a serious human health concern for the foreseeable future and there is an urgent need for the development of therapeutics capable of treating patients and those at high risk for infection and/or with a poor prognosis.

Several lines of evidence suggest that SARS-CoV-2 neutralizing antibodies (nAbs) that bind directly to the virus spike glycoprotein and inhibit entry into host cells have therapeutic potential. First, many infected individuals either remain asymptomatic or recover rapidly with only minimal symptoms, and the plasma from these convalescent patients usually contains nAbs (Long et al., 2020; Robbiani et al., 2020). Second, transfer of plasma containing nAbs from convalescent patients to symptomatic patients has been beneficial in some cases (Duan et al., 2020; Li et al., 2020; Shen et al., 2020). Third, recombinant nAbs that inhibit the interaction between SARS-CoV-2 and host cells confer protection in cell-based assays and animal models (Alsoussi et al., 2020; Shi et al., 2020), and efficacy has also been observed for nAbs targeting the related coronaviruses SARS-CoV (Meulen et al., 2004; Sui et al., 2004; Zhu et al., 2007) and MERS (Corti et al., 2015). Consequently, a number of nAbs have entered clinical trials as post-infection treatment of COVID-19 associated with SARS-CoV-2 (Clinicaltrials.gov - NCT04452318, NCT04497987), with some (Bamlanivimab, Casirivimab, and Imdevimab) receiving Emergency Use Authorization for treatment of mildly ill subjects.

SARS-CoV-2 virions contain 25-100 glycosylated spike (S) proteins that protrude from the viral membrane (Ke et al., 2020; Klein et al., 2020). The S-protein binds to the host cell protein, angiotensin-converting enzyme 2 (ACE2), to mediate viral entry (Hoffmann et al., 2020). The S-protein is a homotrimer and each of its three receptor binding domains (RBD) can be found in either the “up” or the “down” conformation, the former required for ACE2 binding. The most potent nAbs against both SARS-CoV-2 and SARS-CoV bind to the RBD and sterically block its interaction with ACE2 (Cao et al., 2020b; Hansen et al., 2020; Pinto et al.; Rogers et al., 2020; Wan et al., 2020). Consequently, we focused our efforts on developing nAbs that bound to the RBD and competed with ACE2.

To date, all clinically advanced candidate nAbs against SARS-CoV-2 infection have been derived by cloning from B cells of recovered COVID-19 patients or from other natural sources (Cao et al., 2020b; Hansen et al., 2020; Noy-Porat et al., 2020; Rogers et al., 2020; Shi et al., 2020; Wan et al., 2020; Wec et al., 2020). Here, we applied an alternative strategy using *in vitro* selection with phage-displayed libraries of synthetic Abs built on a single human IgG framework derived from a clinically validated drug, trastuzumab (Cobleigh et al., 1999). This approach enabled the rapid production of high affinity nAbs with drug-like properties ready for pre-clinical assessment. Moreover, the use of a highly stable framework enabled facile and modular design of ultra-high-affinity nAbs in tetravalent formats that retained favorable drug-like properties while exhibiting neutralization potencies that greatly exceeded those of the bivalent IgG format. Our tetravalent platform provides a general approach for rapidly improving the potency of virtually any nAb targeting pathogen-related receptor binding proteins, including SARS-CoV-2 and related coronaviruses. Thus, our strategy can improve existing COVID-19 nAb drugs and can be adapted in response to resistant mutations or to future viral threats.

## RESULTS

### Engineering of anti-RBD Fabs and IgGs

Using a phage-displayed human antigen-binding fragment (Fab) library similar to the highly validated library F (Persson et al., 2013), we performed four rounds of selection for binding to the biotinylated RBD of the S-protein of SARS-CoV-2 immobilized on streptavidin-coated plates. Screening of 384 Fab-phage clones revealed 348 that bound to the RBD but not to streptavidin. The Fab-phage were screened by ELISA and those exhibiting >50% loss of binding to the RBD in the presence of 200 nM ACE2 were sequenced, resulting in 34 unique clones (**Fig. 1A**) that were converted into the full-length human IgG1 format for purification and functional characterization.

**Figure 1.**
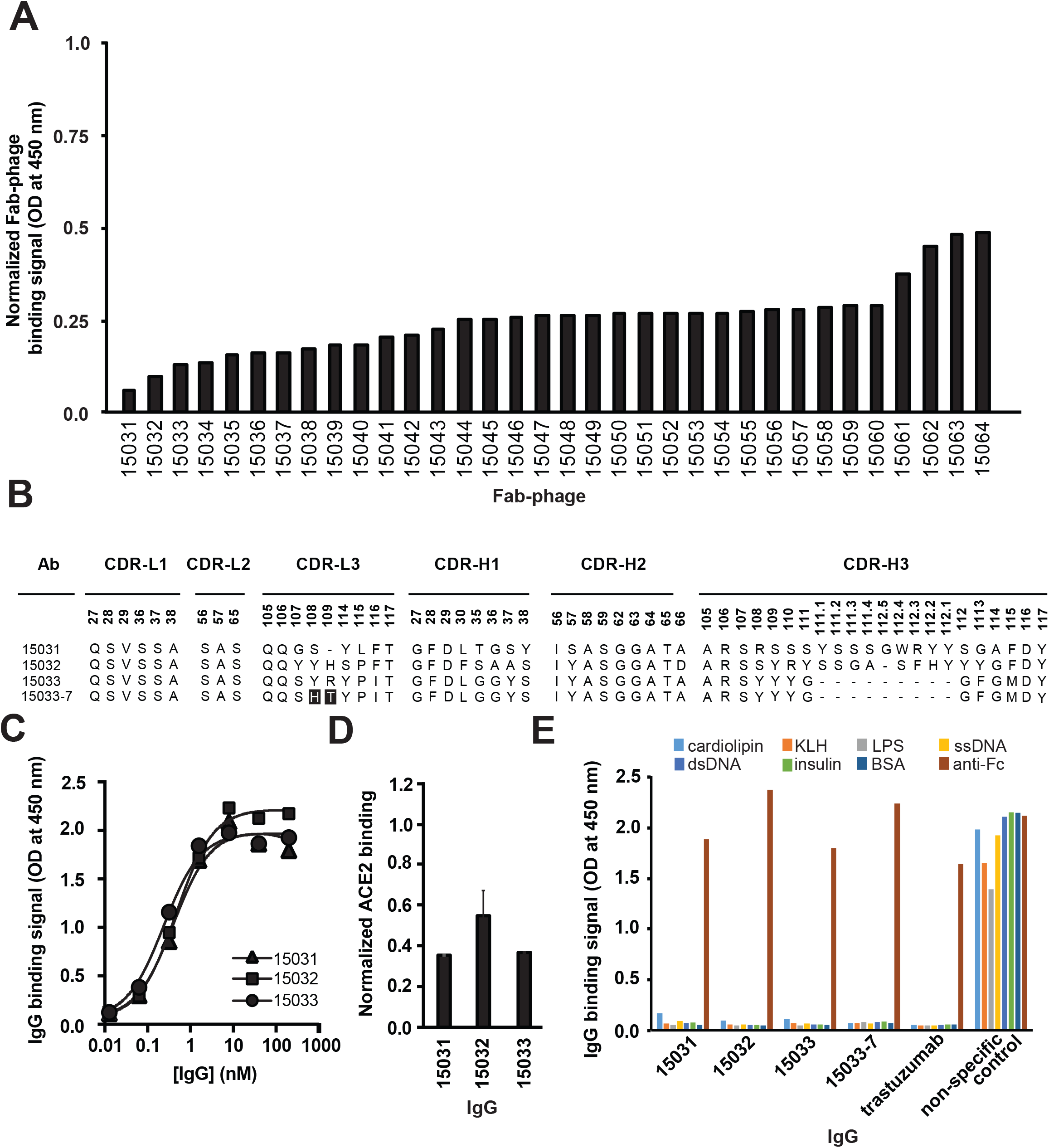
Characterization of anti-RBD Abs by ELISA. **(A)** Binding of unique Fab-phage clones to immobilized RBD blocked by solution-phase ACE2. Signal was normalized to the signal in the absence of ACE2. **(B)** CDR sequences of Abs for which the binding to RBD was strongly blocked by ACE2. Positions are numbered according to the IMGT nomenclature (Lefranc et al., 2003). Sequences in 15033-7 that differ from 15033 are shaded *black.* **(C)** Serial dilutions of IgG binding to immobilized S-protein trimer. The EC_50_ values derived from the curves are shown in **Table 1** and values are representative of 2 independent experiments. **(D)** Binding of biotinylated ACE2 to immobilized S-protein blocked by solution-phase IgG. Signal was normalized to the signal in the presence of a non-binding control IgG and error bars show the standard error of the mean of duplicate samples. (**E**) Assessment of non-specific binding of IgGs to immobilized antigens or a goat anti-human Fc Ab (positive control).

To determine relative binding strength, ELISAs were performed with serial dilutions of IgG protein binding to biotinylated S-protein trimer captured with immobilized streptavidin. These assays showed that three IgGs bound with EC_50_ values in the sub-nanomolar range (**Fig. 1B, C** **and** **Table 1**). Each IgG also partially blocked the binding of biotinylated ACE2 to immobilized S-protein (**Fig. 1D**). Moreover, similar to the highly specific IgG trastuzumab, these three IgGs did not bind to seven immobilized, heterologous proteins that are known to exhibit high non-specific binding to some IgGs. The observed lack of binding to these heterologous proteins is a predictor of good pharmacokinetics *in vivo* (**Fig. 1E**) (Jain et al., 2017; Mouquet et al., 2010). We also used biolayer interferometry (BLI) to measure binding kinetics and to determine avidities more accurately. All three antibodies exhibited sub-nanomolar K_D_ apparent (**Table 1**), in agreement with the estimates determined by ELISA (**Fig. 1C**). Among these, IgG 15033 exhibited the highest avidity, which was mainly due to a two- or seven-fold higher on-rate than IgG 15031 or 15032, respectively. Based on the binding kinetics, we focused further efforts on Ab 15033.

**Table 1.**
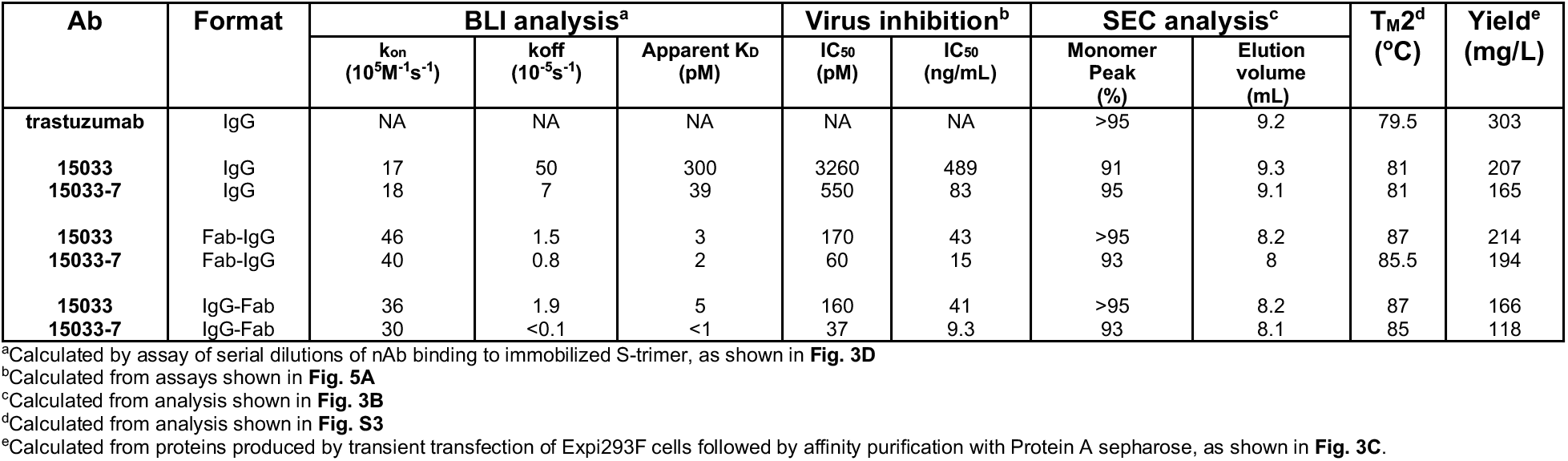
Affinity, potency and biophysical characteristics of nAbs.

We took advantage of the modular design principles of our synthetic Ab library to improve the affinity of Ab 15033. The naïve synthetic library was constructed with tailored diversification of key positions in all three heavy chain complementarity-determining regions (CDRs) and the third CDR of the light chain (CDR-L3). We reasoned that the already high affinity of Ab 15033 could be further improved by recombining the heavy chain with a library of light chains with naïve diversity in CDR-L3. Following selection for binding to the RBD, the light chain library yielded numerous variants, of which 17 were purified in the IgG format and analyzed by BLI. Several of the variant light chains resulted in IgGs with improved binding kinetics compared with IgG 15033, and in particular, IgG 15033-7 (**Fig. 1B**) exhibited significantly improved avidity (K_D_ apparent = 300 pM for 15033 or 39 pM for 15033-7) due to an off-rate that was an order of magnitude slower (**Table 1**).

### Structural analysis of Fabs in complex with the RBD and the S-protein

To rationalize the molecular basis for the differences between nAbs 15033 and 15033-7, and their ability to block ACE2 binding, we first solved the X-ray crystal structures of Fabs in complex with the SARS-CoV-2 RBD at 3.2 and 3.0 Å resolution, respectively (**Fig. 2A**, **Table S1**). As expected, backbone superposition showed that the two complexes were essentially identical (RMSD = 0.17 Å). The binding of Fab 15033-7 to the RBD resulted in an interface with 1130 and 1112 Å^2^ of buried surface area on the RBD and Fab, respectively. Of the surface area buried on the Fab, 59% comes from the light chain and 41% from the heavy chain, with the Fab paratope centered on the receptor binding motif of the RBD (**Fig. 2B**). Comparison of the Fab and ACE2 footprints on the RBD revealed that they overlap extensively, with 69% of the ACE2 footprint covered by that of the Fab footprint (**Fig. 2C**). It follows that direct steric hindrance explains the ability of 15033 and 15033-7 to block the RBD-ACE2 interaction (**Fig. 1D**).

**Figure 2.**
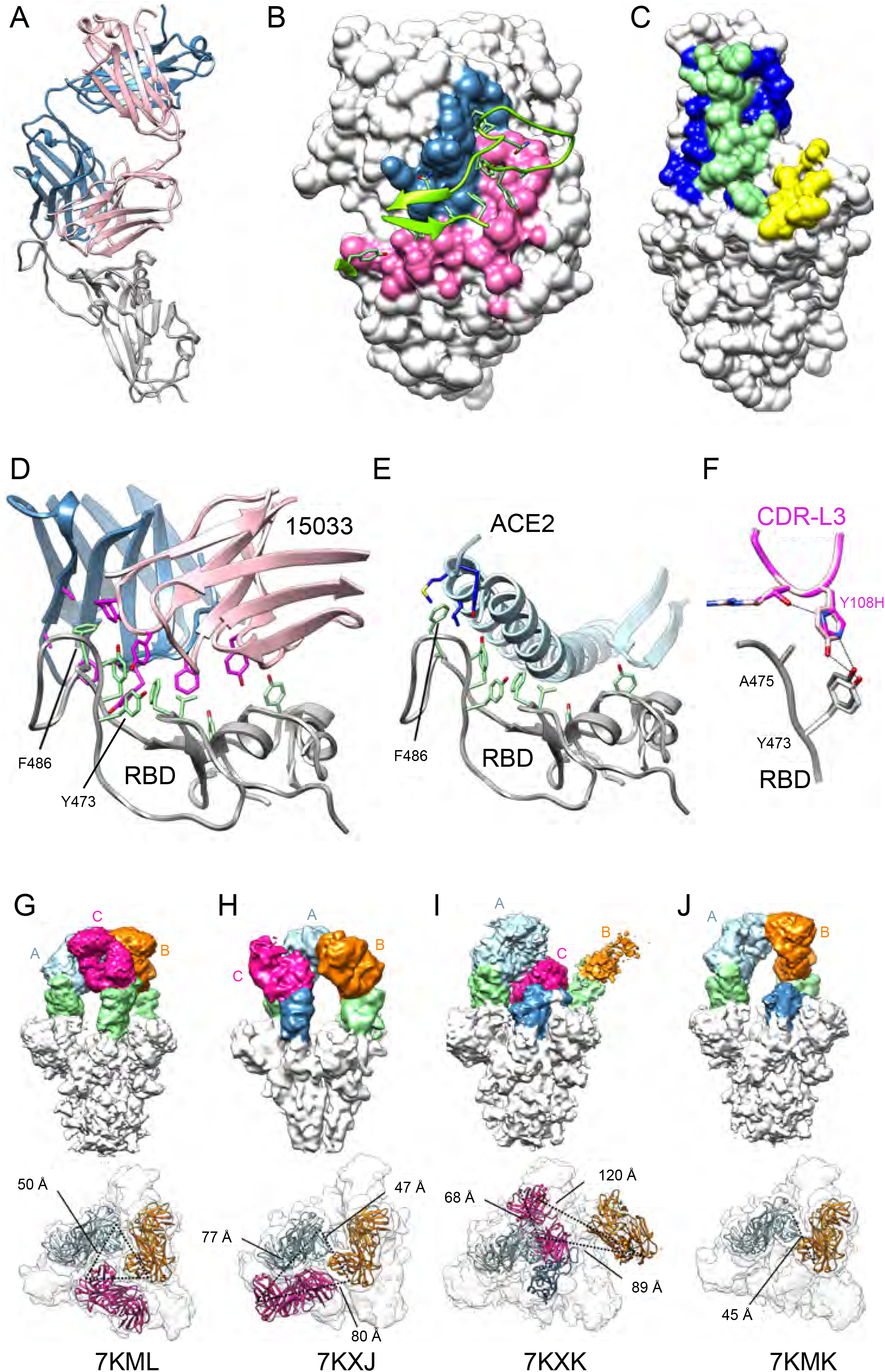
Structural analysis of Fabs bound to the RBD. **(A-F)** X-ray crystallography analysis. **(A)** Ribbon diagram of Fab 15033-7 in complex with the RBD. The RBD is shown in white. The light and heavy chains of the Fab are shown in pink or blue, respectively. **(B)** Fab 15033-7 (surface representation) and its interaction with the receptor binding motif (green ribbons) of the RBD. The paratope defined by the Fab-RBD complex is colored blue (heavy chain) and pink (light chain). **(C)** Surface representation of the RBD colored to show the Fab footprint (blue + green), the ACE2 footprint (yellow + green) and their significant overlap (green). **(D)** Non-polar residues (magenta) on the Fab (15033 shown; light chain, pink; heavy chain, blue) make key interactions with a stretch of exposed apolar residues (green) on the RBD (white). **(E)** These same non-polar residues are critical for ACE2 binding. Highlighted is the important interaction with RBD residue Phe^486^ in the ACE2 complex. **(F)** Fabs 15033 (light pink) and 15033-7 (dark pink) differ by only two residues in CDR-L3. In 15033-7, His^108^ forms an additional hydrogen bond with Thr^109^, stabilizing the local conformation of the CDR-L3 loop. **(G-J)** Cryo-EM analysis. Four Fab-S-protein complexes were observed in the cryo-EM maps: **(G)** the 3-Fab-bound, 3-“up”, C3 symmetric structure; **(H)** the 3-Fab-bound, 3-“up”, asymmetric structure; **I)** the 3-Fab-bound, 2-“up”-1-“down” structure; **(J)** the 2-Fab-bound, 2-“up” structure. Upper row: side view of the EM maps; lower row: top view of the EM maps with the Fabs in ribbons. The dotted lines indicate distances between the C-terminal ends of the Fab heavy chains. Light blue, unit A Fab; orange, unit B Fab; magenta, unit C Fab; green/blue, RBD. In **(I)** and **(J)** the blue RBDs are in the “down” position. In **(H)** the blue RBD and its bound Fab (unit C) is rotated, breaking the C3-symmetry. In **(I)** the density for the B unit Fab is weak. The distances involving it were derived from the best fit of a Fab (orange) to the density (also see **Fig S1, S2**).

Fabs 15033 and 15033-7 recognize a patch of surface-exposed non-polar residues on the RBD using several non-polar residues in both their heavy and light chain CDRs (**Fig. 2D**). The significance of this interaction mode is made clear by a comparison with the ACE2-RBD complex (**Fig. 2E**). The same patch of non-polar RBD residues mediates the interaction with ACE2, a reflection of their importance in both the Fab-RBD and ACE2-RBD complexes. The RBD of SARS-CoV-2 binds to ACE2 with high affinity and exploiting this surface on the RBD may explain why Ab 15033 emerged as the most potent Ab in our initial phage-display screen. RBD residue Phe^486^ is particularly noteworthy as its side chain is completely buried in a pocket between CDRs H2, H3 and L3.

Abs 15033 and 15033-7 differ at only two positions (15033, Tyr108L/Arg109L; 15033-7, His108L/Thr109L). In the RBD-15033 complex, Tyr^108L^ makes van der Waals interactions with RBD residues Tyr^473^ and Ala^475^, as well as a weak hydrogen bond to the side chain of Tyr^473^ (**Fig. 2F**). In the higher affinity RBD-15033-7 complex, the side chain of the equivalent residue, His^108L^, makes similar interactions. However, its side chain also makes an additional intramolecular hydrogen bond to the corresponding residue, Thr^109L^, the other residue that differs between 15033 and 15033-7. Taken together, these residue changes likely stabilize the 15033-7 CDR-L3 loop conformation with a concomitant improvement in both the van der Waals and hydrogen bond interactions with RBD residues Tyr^473^ and Ala^475^.

Using electron cryo-microscopy (cryo-EM), we also determined the structure of Fab 15033-7 in complex with the S-protein trimer. This analysis resulted in the identification of at least 4 different complexes with either two or three Fabs bound to each S-protein trimer (**Fig. 2G-J**, **S1, S2**). For all but one of the Fabs in these complexes, interpretable density for the entire Fab was observed, a result of Fab-Fab interactions that served to immobilize the Fab and the RBD to which it was bound.

In two of the complexes with three Fabs bound (**Fig. 2G-H**), all three RBDs were found in the “up” conformation. The Fabs were relatively well-ordered in both complexes, as adjacent Fabs contacted each other around the three-fold rotation axis describing the S-protein trimer. Although three-fold symmetry was observed in one of the complexes (**Fig. 2G**, **S1A**), in the other, one of the Fab-RBD units made unique Fab-Fab interactions that broke the symmetry (**Fig. 2H**, **S1B**).

In the third complex with three Fabs bound, two of the RBDs were in the “up” conformation and the third RBD was in the “down” conformation (**Fig. 2I**, **S1C**). To accommodate the latter, one of the other Fab-RBD units was pushed away from the 3-fold rotation axis where it was unable to make Fab-Fab contacts. As such, the density for the Fab was relatively weak, an indication of motion/disorder. The remaining Fab-RBD unit stacked over the one in the “down” conformation, making extensive and yet again different Fab-Fab contacts. This led to a well-packed arrangement where the Fab bound to the “down” RBD was sandwiched between the two “up” RBDs.

In the complex with 2 Fabs bound (**Fig. 2J**, **S1D**), the bound RBDs were in the “up” conformation and the Fab-Fab interactions were very similar to those found in the symmetrical complex with three Fabs bound (**Fig. 2G**, **S1A**). The structure showed that the unbound RBD, which was in the “down” conformation, could not sterically accommodate a Fab and further affirmed that one of the two “up” Fab-RBD units must move away from the 3-fold rotation axis if binding to it was to occur.

Analysis of the distances between the C-termini of the heavy chains of the Fabs in these complexes showed that they range from 45 to 120 Å (**Fig. 2G-J**, lower row). In some cases, they are oriented such that two of them, with some repositioning of the Fab-RBD unit, could be linked to a single IgG molecule. Indeed, as described below, negative stain electron microscopy confirmed simultaneous binding of both Fab arms of a single IgG 15033-7 molecule to the S-protein trimer.

Taken together, our structural analysis showed that Fab 15033/15033-7 blocks ACE2 binding to the RBD by direct steric hindrance and that the simultaneous binding of both IgG arms to the S-protein trimer likely enhances potency through avidity effects. It also showed that Fab 15033/15033-7 can make four conformationally distinct complexes with the S-protein trimer through interactions involving the RBD in either the “up” or “down” conformation, a property also shared by some other ACE2-blocking Fabs (Barnes et al., 2020). While it was not immediately apparent why these Fabs were able to recognize both “up” and “down” positions, such flexibility bodes well for efforts aimed at the design of improved antibody-based therapeutics. Finally, it showed that different Fab-Fab interaction modes served to define and stabilize the complexes observed.

### Engineering of tetravalent nAbs with enhanced avidities

Building on these observations, we explored whether we could further enhance the interactions of these nAbs with the S-protein trimer by taking advantage of modular design strategies. We generated tetravalent versions of 15033 and 15033-7 by fusing additional copies of the Fab to either the N- or C- terminus of the IgG heavy chain to construct molecules that we termed Fab-IgG or IgG-Fab, respectively (**Fig. 3A**).

**Figure 3.**
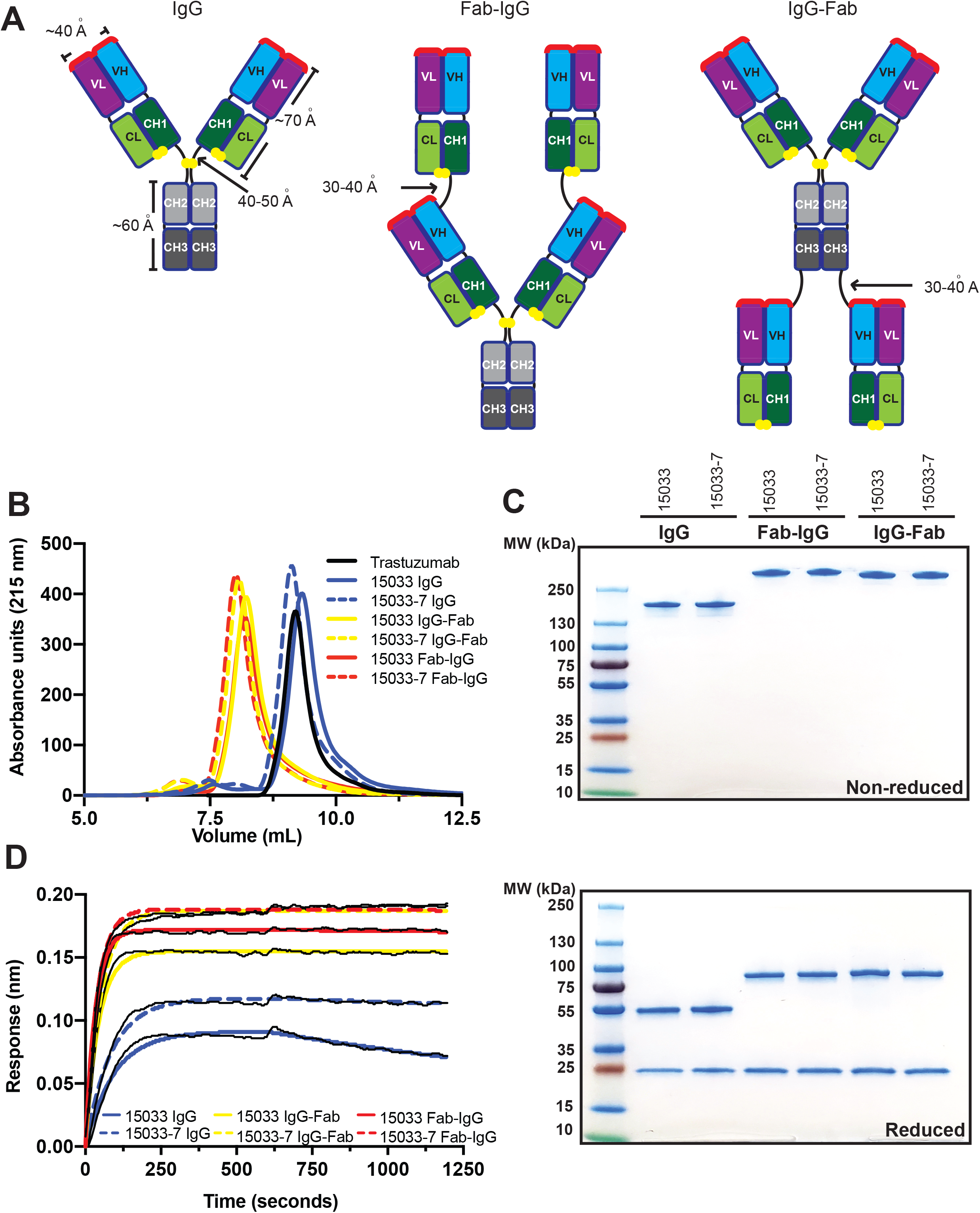
Design and characterization of tetravalent nAbs. **(A)** Schematic of the bivalent IgG format, and the tetravalent Fab-IgG and IgG-Fab formats. Paratopes are shown in red, linkers are shown in black, and disulfide bonds are shown as yellow spheres. **(B)** Analytical gel filtration SEC of nAbs. **(C)** SDS-PAGE analysis of nAbs under non-reducing (top) or reducing conditions (bottom). **(D)** BLI sensor traces for nAbs (6.7 nM) binding to immobilized S-protein trimer.

Our ultimate goal is to produce nAbs that can be used to counter SARS-CoV-2 infections either as a therapeutic and/or as a prophylactic. Aside from high affinity and specificity, effective nAb drugs must also possess favorable properties including high yields from recombinant expression in mammalian cells, high thermodynamic stability, and the lack of aggregation and excessive hydrophobic surface area. To examine these properties, we produced IgGs 15033 and 15033-7, and their Fab-IgG and IgG-Fab counterparts, by transient expression in Expi293F cells. All six proteins were expressed in high yield (160-200 mg/L) and showed high thermostability with CH3/Fab melting temperatures ranging from 81-87 °C, values which exceeded that of the trastuzumab Fab (79.5 °C, **Table 1**, **Fig. S3**). Size exclusion chromatography revealed that each IgG eluted as a predominant (91 to >95%) monodispersed peak with elution volumes nearly identical to that of trastuzumab (**Fig. 3B** **and** **Table 1**). All the tetravalent molecules eluted as single peaks in advance of trastuzumab, an observation consistent with their larger molecular weights. We also showed that the IgG and tetravalent versions of both 15033 and 15033-7 could be purified to near homogeneity by protein-A affinity chromatography as evidenced by SDS-PAGE (**Fig. 3C**). Taken together, these analyses demonstrated that the IgGs and their tetravalent derivatives possess excellent biophysical properties that will facilitate drug development and production at large scale.

Importantly, the tetravalent Abs exhibited greatly reduced off-rates compared with their bivalent IgG counterparts, with apparent dissociation constants for the S-protein trimer in the low single-digit picomolar range as measured by BLI (**Fig. 3D**, **Table 1**). Negative stain electron microscopy of IgG 15033-7, in complex with the S-protein trimer, showed that the two arms of a single IgG bound the S-protein trimer in a pincer-like fashion (**Fig. 4B,D**). Fab-IgG 15033-7 also makes pincer-like interactions with a single S-protein trimer **(Fig. 4C,E**, **Fig. S6, S7)**, but the complexes revealed additional density that differed from those observed for the complex of the S-protein trimer with the IgG. In particular, we observed density consistent with that from an additional Fab in the Fab-IgG complexes, suggesting that all three of the S-protein RBDs are bound by Fabs from a single tetravalent nAb molecule, Fab-IgG in this instance. Given the range of complexes observed in our cryo-EM analysis, it is possible that the Fab-IgG, with four available Fabs, can engage all three RBDs within a single S-protein molecule (**Fig. 4E**). Further analysis will be required to fully establish whether this occurs, which could explain the decreased off-rates shown by the tetravalent proteins due to avidity. Nevertheless, it is clear that both the Fab-IgG and IgG-Fab possess an increased potential for multivalent interactions with one or more S-protein trimers. The latter mode of interaction may be possible on the viral surface and likely contributes to enhanced potency observed for the tetravalent IgGs, as shown below using a cell-based infectivity assay.

**Figure 4.**
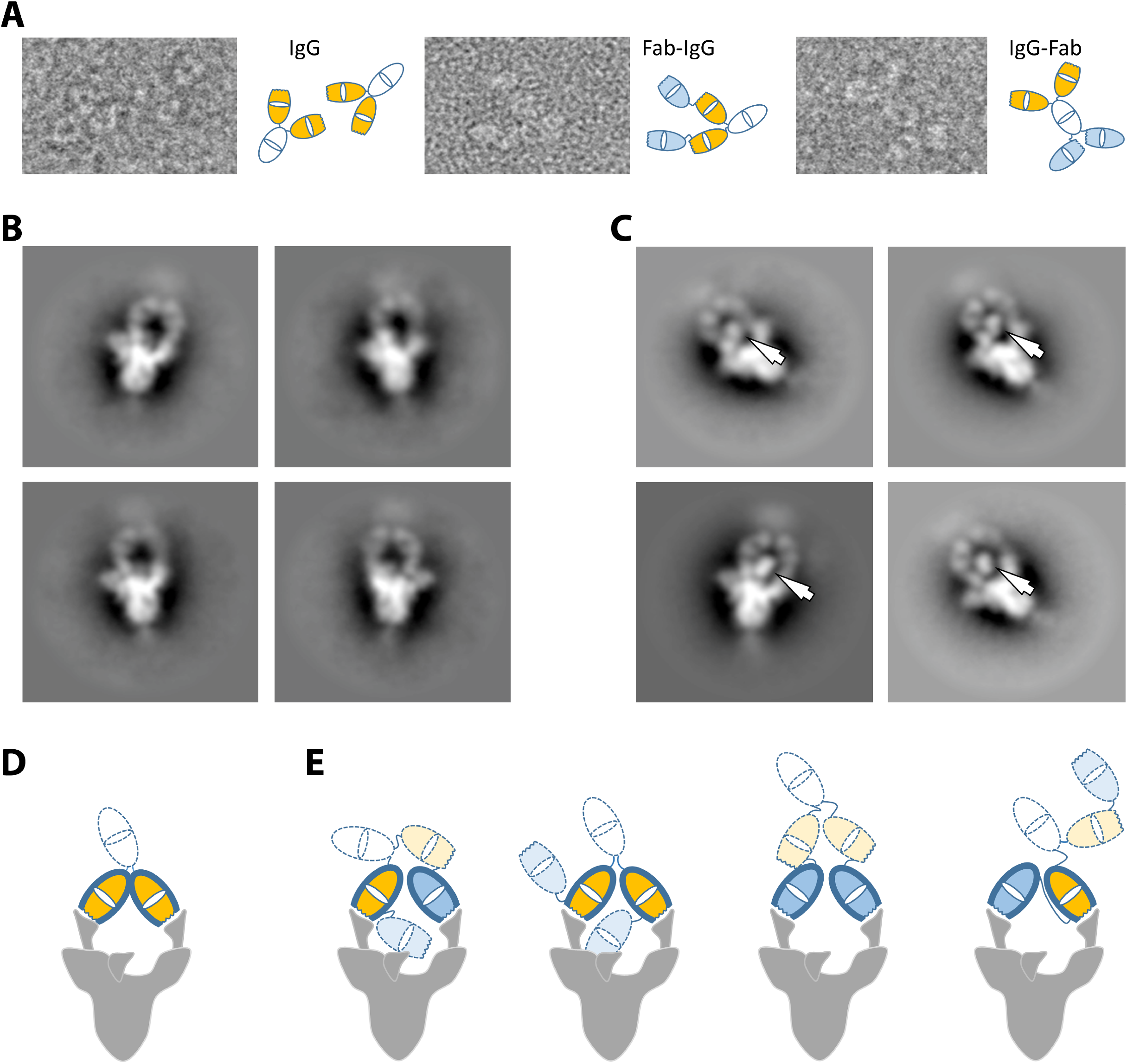
Negative stain electron microscopy analysis of nAbs and their interaction with the S-protein trimer. **(A)** Examples of the 15033-7 IgG (left), Fab-IgG (middle) and IgG-Fab (right) molecules observed in the negative stain micrographs. Schematic interpretations are shown to the right of each image. **(B)** Negative stain electron microscopy 2D class averages of IgG 15033-7 in complex with the S-protein trimer. **(C)** Negative stain electron microscopy 2D class averages of Fab-IgG 15033-7 in complex with the S-protein trimer. The arrows indicate Fab-sized densities not observed in the IgG complex in **(B)**. **(D)** Tentative schematic of the complex in **(B)** showing a trimer with 2 RBDs “up” and one RBD “down”. **(E)** Schematics of the four possible ways that the Fab-IgG can bind to the two “up” RBDs of a trimer that has two RBDs “up” and one RBD “down”. In **(D)** and **(E)**, the following color scheme was used: white, Fc; orange, the Fabs of a native IgG; blue, the Fabs added to the native IgG. Wavy lines indicate the locations of the CDRs. The Fabs with heavy outline are making specific interactions through their CDRs. As shown by our Cryo-EM structures, the RBD in the “down” conformation is accessible for specific interaction with one of the remaining Fabs (orange or blue with light dotted outline) shown in **(E)**. This would lead to a trivalent interaction with the S-protein trimer and may explain the additional Fab-sized densities observed in **(C)**.

### Inhibition of SARS-CoV-2 infection in cell-based assays

We assessed the effects of the nAbs on virus infection in an assay that measured the infection of ACE2-expressing Vero E6 cells with the SARS-CoV-2 strain 2019 n-CoV/USA_WA1/2020. All three high affinity nAbs (15031, 15032 and 15033) from the naïve library (**Fig. 1**) exhibited dose-dependent neutralization of SARS-CoV-2 infection, confirming their inhibitory capacity (**Fig. S4**). Consistent with their affinities, IgG 15033 was the most potent with an IC_50_ of 3.3 nM, and its potency was confirmed with the observation of strong neutralization of a second SARS-CoV-2 strain (2019-nCoV/Italy-INMI1). Neutralization of the second strain was consistent with that of the first strain, and for simplicity, only one set of data are shown.

The tetravalent Fab-IgG and IgG-Fab versions of 15033 exhibited improved potencies with IC_50_ values of 170 and 160 pM, respectively (**Fig. 5A** **and** **Table 1**). The optimized IgG 15033-7 also exhibited high neutralizing potency with an IC_50_ of 550 pM and its tetravalent Fab-IgG and IgG-Fab versions exhibited the best potencies of all the molecules tested with IC_50_ values of 60 and 37 pM, respectively. Taken together, these results support the ability of naïve synthetic Ab libraries to yield highly potent nAbs with drug-like properties. Moreover, further optimization through the engineering of tetravalent formats can produce drug-like molecules with ultra-high potencies that exceed those of bivalent IgGs.

**Figure 5.**
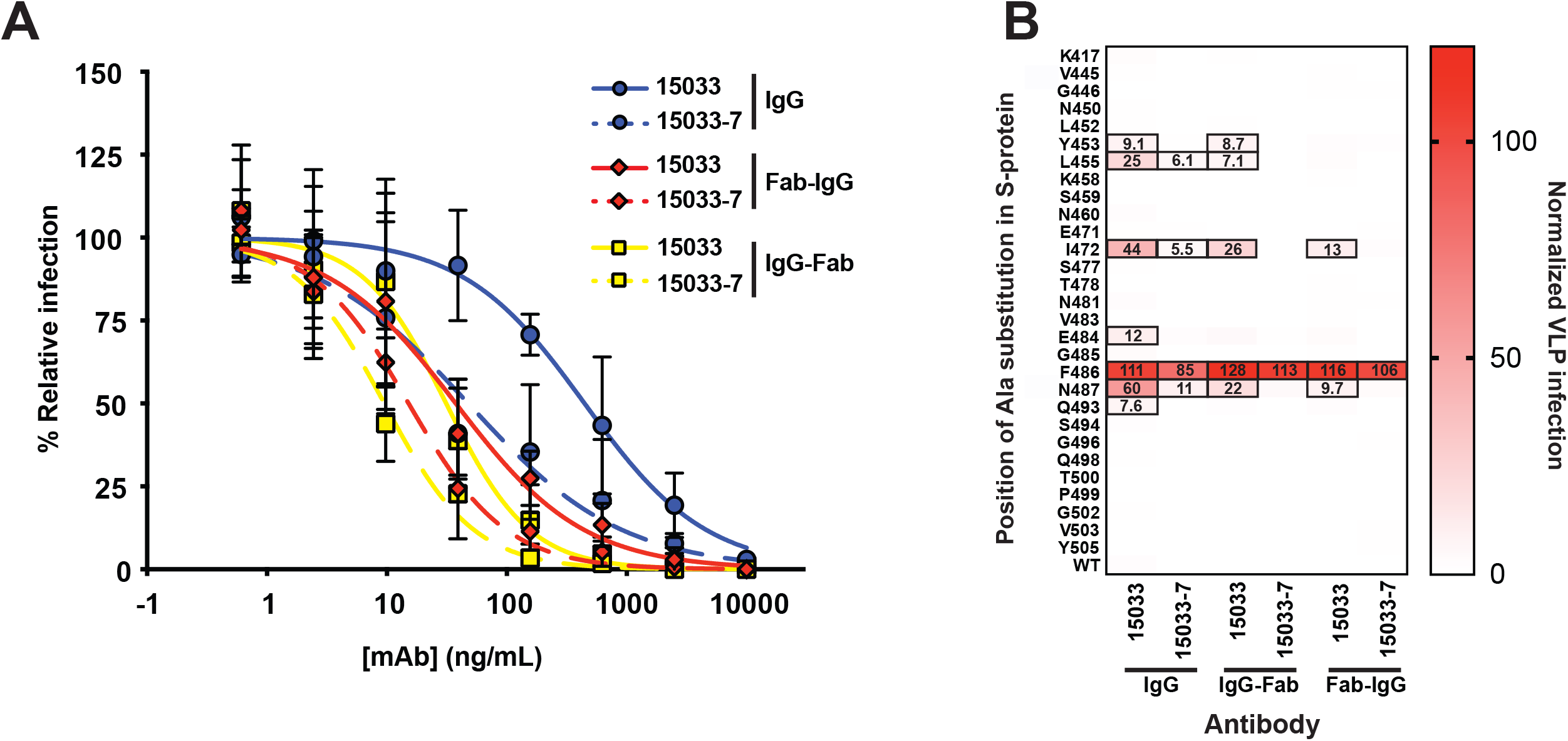
Neutralization of SARS-CoV-2 and pseudotyped VLPs. **(A)** Neutralization of SARS-CoV-2 strain 2019 n-CoV/USA_WA1/2020 by bivalent and tetravalent nAbs (also see **Fig. S4**). The virus was pre-treated with serial dilutions of nAb and infection of ACE2-expressing Vero E6 cells was measured relative to untreated control. Samples were run in triplicate and results are representative of two independent experiments. Error bars indicate standard error of the mean. **(B)** Neutralization of a panel of pseudotyped VLPs displaying SARS-CoV-2 S-proteins with single alanine mutations in or near the ACE2-binding site of the S-protein RBD (also see **Fig. S5**). The VLPs were treated with 50 nM of the indicated nAb and uptake by ACE2-expressing HEK-293 cells was measured in duplicate, and results are representative of two independent experiments. The heat map shows uptake normalized to uptake in the absence of nAb. Boxed cells indicate VLPs that represent potential escape mutants for a given nAb, as defined by >5% uptake with nAb treatment compared with untreated control (the percent uptake is shown in each cell).

### Resistance to potential viral escape mutants

To explore the sensitivity of our nAbs to potential escape mutants, we generated HIV-gag-based lentivirus-like particles (VLPs) pseudotyped with the SARS-CoV-2 S-protein. We confirmed ACE2-dependent infection of HEK-293 cells stably over-expressing exogenous ACE2 by pseudotyped VLPs, and we showed that infection was inhibited by either Fc-tagged RBD (RBD-Fc) or IgG 15033 (data not shown). Using this system, we generated a panel of 44 pseudotyped VLP variants **(Fig. S5A)**, each containing a single alanine substitution at an RBD position within or close to the ACE2-binding site. Twenty of these VLP variants exhibited a >4-fold reduction in infection compared with the wild-type (wt) VLP, suggesting that the substituted residues contributed favorably to the interaction between the RBD and ACE2. The remaining 29 VLP variants infected with high efficiency, suggesting that these are positions where residue changes could abrogate antibody binding without affecting the ACE2 interaction.

Using the panel of 29 infective VLP variants, we measured VLP infection after treatment with the various nAbs (**Fig. 5B**). We defined as escape mutants those VLP variants for which entry in the presence of 50 nM nAb was >5% of the entry in the absence of the nAb. Based on this definition, we found that 7 of the variants were able to escape from IgG 15033 (**Fig. S5A**), whereas only 4 could escape from IgG 15033-7. We also found that 15033 in tetravalent Ab format neutralized more variants than it did as an IgG, and that, remarkably, tetravalent Ab 15033-7 strongly neutralized all of the variants but one (Phe^486^). As discussed above, RBD residue Phe^486^ is found in the non-polar interface between the RBD and the Fab and it makes numerous contacts with CDRs H2, H3 and L3, sitting deep in a pocket formed by these contacts (**Fig. S5B,C**). As such, it represents a bona fide positional vulnerability where mutant viruses still capable of binding ACE2 could evade neutralization by nAb 15033-7. Comprehensive maps of escape mutations confirm the vulnerability of other nAbs to this mutation, but also suggest strategies by which escape can be overcome with nAb cocktails or bi-specific nAbs (Greaney et al., 2020). Overall, these results showed that for IgGs, increasing the affinity of the Fab-RBD interaction increased neutralizing potency and resistance to mutation, and moreover, that tetravalent presentation of the Fab provides even greater potency and increased resistance to potential escape mutants.

## DISCUSSION

SARS-CoV-2 has wreaked havoc on global health and the economy and highlighted the need for drug development technologies to combat not only COVID-19, but emerging infectious diseases in general. In this context, we have used synthetic antibody engineering to rapidly develop unique formats of human nAbs as therapeutic candidates. Our nAb formats include natural bivalent IgGs as well as ultra-potent tetravalent molecules arising from a modular design strategy that adds two additional Fabs to the canonical IgG molecule. Most importantly, we have shown that the enhanced avidities and potencies characteristic of the tetravalent nAbs are achieved without compromising the favorable properties – such as yield, solubility, and stability - that make IgG molecules ideal drugs. Moreover, we showed that the tetravalent nAbs are more resistant to potential escape mutations, an observation highlighting the utility of these molecules as therapeutics against SARS-CoV-2 and other viruses capable of rapidly mutating under selection pressure.

Structural analyses showed that IgG and Fab-IgG 15033-7 make pincer-like interactions with a single SARS-CoV-2 S-protein trimer, an observation providing insight into the basis for the avidity observed for these molecules. It also provides strong evidence that the tetravalent Fab-IgG can engage all three RBDs of the S-protein trimer, and that the RBD can be bound by a Fab in either the “up” or “down” conformation. We did not optimize the tetravalent formats; their very high avidities arose naturally from the geometry of the Fab-RBD complex, the spatial relationships between the Fabs, the flexibility of the RBDs on the S-protein trimer, and fortuitous Fab-Fab interactions that clearly stabilized the complexes observed. We and others have generated many additional Fabs that can inhibit the virus but employ different epitopes to do so, and incorporating these Fabs into tetravalent frameworks, including bispecific formats targeting two distinct epitopes, has the potential to generate a vast array of ultra-high affinity nAbs with minimal further effort. Taken together, these observations underscore the vast potential of our modular design approach for the development of novel and highly effective nAb therapeutics, and our approach has the potential to further optimize any COVID-19 nAb therapeutic currently in development.

COVID-19 has also exposed the need for drug development to respond to emerging viral threats in real time. In this regard, the isolation of nAbs from the B cells of infected individuals has emerged as a rapid approach to obtaining leads for novel drug development. Indeed, several reports have shown that these technologies can deliver drug-grade therapeutic nAbs for manufacturing and subsequent clinical trials in approximately six months (Hansen et al., 2020; Jones et al.). Further highlighting the urgent need and the rapid speed at which drug development has proceeded, these IgG-based nAbs have now received emergency-use approval from the US FDA as both cocktail and single agent, and these extraordinarily rapid drug development timelines set a benchmark for alternative technologies. The potencies of other anti-SARS CoV-2 nAbs obtained in similar fashion range from nanomolar (Wu et al., 2020) to picomolar (Liu et al., 2020), but those that have been awarded approval for emergency use have potencies in the middle of this range (Hansen et al., 2020; Jones et al.). Thus, it is worth noting that antibodies with higher *in vitro* potencies may further improve *in vivo* efficacy, and our tetravalent formats should be applicable to enhance potency of nAbs in general.

With our platform, we show that synthetic *in vitro* antibody engineering is comparable to B cell cloning, in terms of both speed and potency of drug development (Hansen et al., 2020; Zost et al., 2020). Our project was initiated at the beginning of April 2020 when we identified our first naïve Ab leads. Within a month, we validated lead IgG molecules as neutralizing agents in cell-based assays with authentic virus. In parallel, we initiated further selections to optimize the Fab 15033 paratope, that yielded our best lead nAb 15033-7; both Fabs were then reformatted as tetravalent molecules (Fab-IgG and IgG-Fab) to further enhance potency. Though others have now shown enhanced potency of SARS-CoV-2 neutralization with oligomers of Ab variable domains (Schoof et al., 2020) and synthetic proteins (Cao et al., 2020a), insofar as our format closely resembles natural Abs, they can also be manufactured at large scale, possess long half-lives and likely low immunogenicity, all of which are required to become effective drugs. Indeed, by the beginning of October, we established manufacturing-grade stable cells capable of producing multi-gram quantities of drug-grade nAbs from a litre of culture, in both the IgG and tetravalent formats. We are now manufacturing our best nAb for clinical trials. Thus, synthetic Ab engineering technologies can match the six-month lead-to-manufacture timelines established by methods based on the cloning of natural nAbs.

Our synthetic engineering technologies offer exquisite control over Ab design and by introducing tailored diversity into an optimized IgG framework, ensures that candidate therapeutics possess biophysical properties that are ideal for drug development. Facilitated by the use of highly stable frameworks, we now demonstrate the rapid construction of complex tetravalent formats that enhance potency while retaining favorable drug-like properties. Perhaps most importantly though, synthetic methods do not rely on infected patients (i.e. natural repertoires) as a source of drug leads. By its very nature, B cell cloning is a reactive technology that can be implemented only after a viral outbreak is underway, and this places a limit on the time required for drug development. With synthetic *in vitro* methods, drug development can proceed in a proactive manner, as the development and stockpiling of potential therapeutics in advance of outbreaks is feasible. Indeed, large-scale surveillance and sequencing efforts have provided unprecedented access to the genomes of numerous SARS-CoV-2-related viruses and other pathogens with the potential to cross species barriers and infect humans (Daszak et al., 2020; Shi and Hu, 2008). With our approach, it is feasible to develop - in advance, synthetic Abs against hundreds of antigens in parallel (Hornsby et al., 2015) and working within a collaborative international network, we have initiated efforts with this aim.

## ACKNOWLEDGEMENTS

This study was supported in part by contracts and grants from NIH (R01 AI157155) and the Defense Advanced Research Project Agency (HR001117S0019) to MSD, NIH grants P01AI120943 and R01AI123926 to GKA, and an NIH grant R01AI107056 to DWL. This study was also supported in part by a CIHR operating grant (COVID-19 Rapid Research Funding #OV3-170649) to JR and SSS, and by the Lazio Region (Italy, DGR n. 653 29 September 2020) and the Rome Foundation (Italy, Prot 317A/I). Additional support was generously provided by FAST grants #2161 to GKA and SSS and #2189 to SSS, from Emergent Ventures through the Thistledown Foundation (Canada) and the Mercatus Center at George Mason University. We are grateful to Daniele Lapa (INMI), for his contribution in determining the neutralizing power of the antibodies tested in Rome. We are also grateful to Carlo Tomino for his valuable help in preparing for the regulatory aspects of monoclonal antibodies in Italy.

## AUTHOR CONTRIBUTIONS

^**†**^**Shane Miersch** Conceptualization, Methodology, Investigation, Supervision, Writing - Original Draft, Writing - Review & Editing, Visualization, Validation, Project administration

^**†**^**Zhijie Li** Methodology, Investigation, Data Curation, Visualization, Formal analysis, Validation, Writing - Original Draft, Writing - Review & Editing

**Reza Saberianfar** Methodology, Investigation

**Mart Ustav** Conceptualization, Methodology, Investigation

**Brett Case** Investigation, Validation

**Levi Blazer** Conceptualization, Investigation

**Chao Chen** Investigation

**Wei Ye** Investigation

**Alevtina Pavlenco** Investigation

**Maryna Gorelik** Investigation

**Julia Garcia Perez** Investigation

**Suryaseree Subramania** Investigation

**Serena Singh** Investigation

**Lynda Ploder** Investigation

**Safder Ganaie** Investigation

**Daisy Leung** Investigation

**Rita E. Chen** Investigation

**Pier Paolo Pandolfi** Investigation

**Giuseppe Novelli** Investigation

**Giulia Matusali** Investigation

**Francesca Colavita** Investigation

**Maria R. Capobianchi** Investigation

**Suresh Jain** Funding acquisition

**J.B. Gupta** Funding acquisition

**Gaya Amarasinghe** Supervision, Writing - Review & Editing

**Michael Diamond** Supervision, Writing - Review & Editing

^**\***^**James Rini** Conceptualization, Writing - Review & Editing, Supervision, Funding acquisition

^**\***^**§Sachdev Sidhu** Conceptualization, Writing - Original Draft, Supervision, Funding acquisition

## STAR METHODS

### Cells

Mammalian cells were maintained in humidified environments at 37 °C in 5% CO_2_ in the indicated media. Vero E6 (ATCC), HEK293T (ATCC) and HEK293T cells stably overexpressing ACE2 were maintained at 37 °C in 5% CO_2_ in DMEM containing 10% (vol/vol) FBS. Expi293F cells (ThermoFisher) were maintained at 37 °C in 8% CO_2_ in Expi293F expression media (ThermoFisher).

### Protein production

The previously reported piggyBac transposase-based expression plasmid PB-T-PAF (Li et al., 2013) containing a CMV promotor (PB-CMV) and a woodchuck hepatitis virus posttranscriptional regulatory element (WPRE) was used for large-scale transient expression. cDNA encoding the SARS-CoV-2 S-protein ectodomain trimer (residues 1-1211), followed by a foldon trimerization motif (Tao et al., 1997), a 6xHis tag and an AviTag biotinylation motif (Fairhead and Howarth, 2015) was cloned in to the PB-CMV vector using standard molecular biology techniques, and residues 682–685 (RRAR) and 986–987 (KV) were mutated to SSAS or PP sequences, respectively, to remove the furin cleavage site on the SARS-CoV-2 spike protein or to stabilize the pre-fusion form of the spike (Pallesen et al., 2017), respectively. The SARS-CoV-2 receptor binding domain (RBD, residues 328-528), the soluble human ACE2 construct (residues 19-615), and the SARS-CoV RBD (residues 315-514), each followed by a 6xHis tag and an AviTag, were similarly cloned into the same vector. For expression, PB-CMV expression constructs were mixed with Opti-MEM media (Gibco) containing 293fectin reagent (Thermo Fisher) and the mixture was incubated for 5 min before addition to the shaker flask containing 10^6^ Freestyle 293-F cells/mL grown in suspension in Freestyle 293 expression media (Thermo Fisher). Expression was allowed to continue for 6 days before purification.

### Protein purification and *in vitro* biotinylation

Expressed proteins were harvested from expression medium by binding to Ni-NTA affinity resin followed by elution with 1X PBS containing 300 mM imidazole and 0.1% (v/v) protease inhibitor cocktail (Sigma), then further purified by size-exclusion chromatography. For the RBDs and ACE2, a Superdex 200 Increase (GE healthcare) column was used. For the S-protein ectodomain, a Superose 6 Increase (GE healthcare) column was used. Purified proteins were site-specifically biotinylated in a reaction with 200 μM biotin, 500 μM ATP, 500 μM MgCl_2_, 30 μg/mL BirA, 0.1% (v/v) protease inhibitor cocktail and not more than 100 μM of the protein-AviTag substrate. The reactions were incubated at 30 °C for 2 hours and biotinylated proteins were then purified by size-exclusion chromatography.

### Phage display selections

A synthetic, phage-displayed antibody library (Persson et al., 2013) was selected for binding to SARS-CoV-2 RBD in solution. In each round, phage library was first depleted on neutravidin immobilized in wells of a 96-well Maxisorp plate from a 2 μg/mL solution incubated with shaking overnight at 4 °C, then incubated with 50 nM biotinylated RBD in solution for two hours at RT. Protein-phage complexes were captured in wells coated with neutravidin as above for 15 min at RT. After washing with 1X PBS pH 7.4 containing 0.05% Tween, phage were eluted for 5 min with 0.1 M HCl and then neutralized with 1 M Tris pH 8.0. Eluted phage were amplified, purified, and selected for binding to target for a total of 5 rounds, after which, individual phage clones were subjected to DNA sequencing, as described (Persson et al., 2013).

### Enzyme-linked immunosorbent assays

For ELISAs, plates were coated with neutravidin, as above, then blocked with PBS, 0.2% BSA for 1 h. Biotinylated target protein was captured from solution by incubation in neutravidin-coated and BSA-blocked wells for 15 min with shaking at RT, and subsequently, phage or Ab was added and allowed to bind for 30 min. Plates were washed, incubated with an appropriate secondary antibody, and developed with TMB substrate as described (Miersch et al., 2017).

### Construction of genes encoding tetravalent Abs

DNA fragments encoding heavy chain Fab regions (VH-CH1; terminating at hinge residue Thr^10^, IMGT numbering (Lefranc et al., 2003)) were amplified by PCR from the IgG expression constructs. Tetravalent Ab constructs were generated by fusing these fragments with their respective IgG heavy chain in the pSCSTa mammalian expression vector using Gibson assembly (New England Biolabs, Ipswich, MA). Fab-IgG constructs were arranged by fusing a heavy chain Fab domain to the N-terminus of the IgG using a S(G4S)_3_ linker. IgG-Fab constructs were arranged by fusing a heavy chain Fab domain to the C-terminus using a G(G4S)_2_GGGTG linker. For both formats, the Fc region terminated at Gly^129^ (IMGT numbering (Lefranc et al., 2003)).

### Ab production and purification

IgG and tetravalent Abs were produced in Expi293F cells (ThermoFisher) by transient transfection, by diluting heavy and light chain construct DNA in OptiMem serum-free media (Gibco) before the addition of and incubation with FectoPro (Polyplus Transfection) for 10 min. For IgG expression, equivalent amounts of plasmids encoding heavy chain or light chain were transfected, whereas for tetravalent formats, a ratio of 2:1 light chain to heavy chain plasmids was used. Following addition of the DNA complex to Expi293F cells and a 5-day expression period, Abs were purified using rProtein A Sepharose (GE Healthcare), then buffer exchanged and concentrated using Amicon Ultra-15 Centrifugal Filter devices (Millipore). IgGs were stored in PBS (Gibco), and tetravalent Abs were stored in 10 mM L-Histidine, 0.9% sucrose, 140 mM NaCl, pH 6.0.

### Size exclusion chromatography

Protein samples (50 μg) were injected onto a TSKgel BioAssist G3SWxl column (Tosoh) fitted with a guard column using an NGC chromatography system and a C96 autosampler (Biorad). The column was preequilibrated in a PBS mobile phase and protein retention was monitored by absorbance at 215 nm during a 1.5 CV isocratic elution in PBS.

### Biolayer interferometry

The binding kinetics and estimation of apparent affinity (KD) of Abs binding to the S-protein were determined by BLI with an Octet HTX instrument (ForteBio) at 1000 rpm and 25 °C. Biotinylated S-protein was first captured on streptavidin biosensors from a 2 μg/mL solution to achieve a binding response of 0.4-0.6 nm and unoccupied sites were quenched with 100 μg/mL biotin. Abs were diluted with assay buffer (PBS, 1% BSA, 0.05% Tween 20) and 67 nM of an unrelated biotinylated protein of similar size was used as negative control. Following equilibration with assay buffer, loaded biosensors were dipped for 600 s into wells containing 3-fold serial dilutions of each Ab starting at 67 nM, and subsequently, were transferred back into assay buffer for 600 s. Binding response data were corrected by subtraction of response from a reference and were fitted with a 1:1 binding model using ForteBio’s Octet Systems software 9.0.

### Differential scanning fluorimetry

Thermostabilities of Abs were determined by differential scanning fluorimetry using Sypro Orange, as described (Niedziela-Majka et al., 2015), with a 1 μM solution of Ab and temperature range from 25-100 °C in 0.5 °C increments.

### Generation of pseudotyped VLPs

HEK-293 cells (ATCC) were seeded in a 6-well plate at 0.3 × 10^6^ cells/well in DMEM (ThermoFisher) supplemented with 10% FBS and 1% penicillin-streptomycin (Gibco) and grown overnight at 37 °C with 5% CO_2_. HEK-293 cells were then co-transfected with 1 μg of pNL4-3.luc.R-E- plasmid (luciferase expressing HIV-1 with defective envelop protein) (NIH AIDS Reagent Program) and 0.06 μg of CMV-promoter driven plasmid encoding wt or mutant S-protein using Lipofectamine™ 2000 transfection reagent (ThermoFisher). Pseudotyped VLPs were harvested by collecting supernatant 48 h after transfection and were filter sterilized (0.44 μm, Millipore Sigma, Cat. No. SLHA033SS).

### Infection assays with pseudotyped VLPs

HEK-293 cells stably over-expressing full-length human ACE2 protein were seeded in 96-well white polystyrene microplates (Corning, Cat. No. CLS3610) at 0.03 × 10^6^ cells/well in DMEM (10% FBS and 1% penicillin-streptomycin) and were grown overnight at 37 °C with 5% CO_2_. Pseudotyped VLPs were mixed with Ab, incubated at room temperature for 10 min, and added to the cells. The cells were incubated at 37 °C with 5% CO_2_, the medium was replaced with fresh DMEM (10% FBS and 1% penicillin-streptomycin) after 6 h, and again every 24 h up to 72 h. To measure the luciferase signal (VLP entry), DMEM was removed and cells were replaced in DPBS (ThermoFisher) and mixed with an equal volume of ONE-Glo™ EX Luciferase Assay System (Promega). Relative luciferase units were measured using a BioTek Synergy Neo plate reader (BioTek Instruments Inc.). The data were analyzed by GraphPad Prism Version 8.4.3 (GraphPad Software, LLC).

### SARS-CoV-2 focus reduction neutralization assay

SARS-CoV-2 strain 2019 n-CoV/USA_WA1/2020 was obtained from the Centers for Disease Control (USA) and Prevention. Virus stocks were produced in Vero CCL81 cells (ATCC) and titrated by focus-forming assay on Vero E6 cells (Case et al., 2020). Serial dilutions of mAbs were incubated with 10^2^ focus-forming units (FFU) of SARS-CoV-2 for 1 h at 37 °C. MAb-virus complexes were added to Vero E6 cell monolayers in 96-well plates and incubated at 37 °C for 1 h. Cells were overlaid with 1% (w/v) methylcellulose in MEM supplemented with 2% FBS. Plates were harvested after 30 h by removing overlays and were fixed with 4% PFA in PBS for 20 min at RT. Plates were washed and sequentially incubated with 1 μg/mL of CR3022 (Yuan et al., 2020) anti-S-protein antibody and HRP-conjugated goat anti-human IgG in PBS, 0.1% saponin, 0.1% BSA. SARS-CoV-2-infected cell foci were visualized using TrueBlue peroxidase substrate (KPL) and were quantitated on an ImmunoSpot microanalyzer (Cellular Technologies). Data were processed using Prism software (GraphPad Prism 8.0).

### X-ray crystallography

The SARS-CoV-2 RBD (residues 328-528) was expressed with a C-terminal 6xHis tag from HEK293F GnT1-minus cells. The Fabs were expressed from HEK293F cells. The RBD was purified from the expression media by metal-affinity chromatography using Ni-NTA beads (Qiagen). The Fab fragments were purified from the expression media using rProtein A Sepharose Fast Flow beads (GE healthcare). The RBD was treated with endoglycosidase H (Robbins et al., 1984) and Carboxypeptidase A (Sigma-Aldrich), to remove the N-glycans and the C-terminal 6xHis tag. The RBD and Fabs were further purified by ion-exchange and hydrophobic interaction chromatography. Before crystallization, the RBD-Fab complexes were purified using size-exclusion chromatography on a Superdex 200 Increase column (GE Healthcare). For both Fab 15033 and Fab 15033-7, the optimized crystallization conditions contained 1.2-1.6 M (NH_4_)_2_SO_4_ and 8-18% glycerol. X-ray diffraction data were collected at 100 K on beamline 08IB-1 at the Canadian Light Source. The diffraction data were integrated and scaled using the XDS package (Kabsch, 2010). The structures were solved by molecular replacement using the program Phaser (McCoy et al., 2007). A homology model of the VH-VL region was produced by the SysImm Repertoire Builder server (https://sysimm.org/rep_builder)(Schritt et al., 2019). The CDR loops were then deleted from this VH-VL model. The constant (CH1-CL) region search model and the RBD search model were both derived from PDB entry 6W41 (Yuan et al., 2020). The three models were used to solve the structure by molecular replacement. The atomic models were built using Coot (Emsley et al., 2010) and refined with Phenix.refine (Afonine et al., 2012).

### Negative stain electron microscopy

3 μL of the S-protein ectodomain were applied to the surface of a glow-discharged (60 s, 15 mA), carbon-coated copper mesh grid. The grid was washed with water and stained with 3 μL of 2% uranyl formate. Micrographs were acquired on a Talos L120C 120 kV electron microscope equipped with a Ceta 16M CMOS camera. Particle selection and two-dimensional classification were carried out using cryoSPARC v2. Contrast transfer function (CTF) estimation was performed using GCTF (Zhang, 2016).

### Cryo-EM

Four additional substitutions for proline residues (817P/A892P/A899P/A942P) were introduced to the S-2P SARS-CoV-2 S-protein ectodomain to stabilize the prefusion conformation (Hsieh et al., 2020). Fab 15033-7 was mixed with the SARS-CoV-2 spike ectodomain at a 3:1 molar ratio. 3 μL of the Fab-spike mixture containing 0.4 mg/mL total protein were applied to a glow-discharged (60 s, 15 mA) C-flat 2/2 carbon holey grid (CF-2/2-4C, Electron Microscopy Science). Plunge vitrification was performed using a Thermo Fisher Scientific Vitrobot Mark IV instrument. The grids were blotted for 2.5 s at 100% humidity and 277 K before being plunge-frozen in liquid ethane cooled to 90 K. Specimen screening and optimization were performed using a Talos L120C 120 kV electron microscope equipped with a Ceta 16M CMOS camera.

High-resolution data were collected on a Thermo Fisher Scientific Titan Krios G3 300 kV microscope equipped with a Falcon 4 Direct Electron Detector. The data were collected at 75000x nominal magnification, resulting in a calibrated pixel size of 1.03 Å. Each movie was recorded in counting mode with a 10 s exposure and saved in 30 fractions. The total exposure was 38 electrons per Å^2^. The data were collected with a 1.0-2.2 μm defocus setting. A total of ~6431 movies were collected for the final high-resolution dataset.

Full-frame motion and local motion corrections of the cryo-EM movies were performed using implementations of the alignframes_lmbfgs algorithm (Rubinstein and Brubaker, 2015) in cryoSPARC v2 (Punjani et al., 2017). CTF parameters were estimated using GCTF (Zhang, 2016). Particle selection was initially performed using a Gaussian blob picker, then by Topaz neural network picking (Bepler et al., 2019), both in cryoSPARC v2. Two-dimensional classification of the particle images was performed using cryoSPARC v2. The particle images corresponding to Fab-S-protein complexes in the different conformations were separated by performing heterogenous refinement in cryoSPARC v2. Initial map generation and homogenous, heterogenous and non-uniform three-dimensional refinements were performed in cryoSPARC v2. For the map with three RBDs in the up conformation, the three-dimensional refinements were carried out with either C1 or C3 symmetry imposed. The maps generated with C3 symmetry were used for model building. A previously reported SARS-CoV-2 spike structure, PDB entry 6VXX (Walls et al., 2020), and the 15033-7:RBD complex reported here, were first docked into the map using UCSF Chimera (Pettersen et al., 2004) and then manually modified using Coot (Emsley and Cowtan, 2004). The HV-LV and HC1-LC domains of the Fab were docked individually to account for elbow angle differences (SI Figure ZFH). The resulting model was refined with C3 symmetry against the cryo-EM map using Rosetta (Wang et al., 2016) and Phenix (Liebschner et al., 2019). The resulting C3-symmetric model was used as starting model to build the other three structures, which were also refined against the corresponding cryo-EM maps using Rosetta and Phenix. The atomic models were validated using Molprobity (Williams et al., 2018) and the comprehensive validation (cryo-EM) tool in Phenix (Liebschner et al., 2019). To improve the interpretability of the Fab-RBD interaction, local refinement of the Fab-RBD unit was performed in cryoSPARC v2 using a mask encompassing one of the Fab-RBD units in the C3-symmetric structure. Moderate resolution improvement for the Fab-RBD unit was achieved (**Fig. S2H**).

### Data deposition

The crystal structures of the RBD-Fab 15033 and RBD-Fab 15033-7 complexes were deposited in the PDB with accession codes 7KLG and 7KLH, respectively. The S-protein-Fab 15033-7 cryo-EM maps and structures were deposited to EMDB and PDB with the following accession codes: EMD-22925/PDB:7KMK (2 Fab bound, 2-“up”-1-“down”), EMD-22926/PDB:7KML (3 Fab bound, 3-“up”, C3-symmetric), EMD-23064/PDB:7KXJ (3 Fab bound, 3-“up” asymmetric), EMD-23065/PDB:7KXK (3 Fab bound, 2-“up”, 1-“down”).

## SUPPLEMENTAL INFORMATION

**Table S1. X-ray data collection and refinement statistics**

**Table S2. Cryo-EM data collection and image processing**

**Figure S1. 2D class averages and resolution plots of the four 15033-7 Fab-S-protein cryo-EM structures.** The GSFSC curve, selected 2D class averages and the local resolution map are shown for each of the four structures: **A)** the 3-Fab-bound, 3-“up”, C3 symmetric structure; **B)** the 3-Fab-bound, 3-“up”, asymmetric structure; **C)** the 3-Fab-bound, 2-“up”-1-“down” structure; **D)** the 2-Fab-bound, 2-“up” structure.

**Figure S2. Cryo-EM maps showing only Fabs and RBDs in the density. A-D)** The 15033-7 Fabs (colored ribbons) are shown in the cryo-EM maps for each of the four structures: **A)** the 3-Fab-bound, 3-“up”, C3 symmetric structure; **B)** the 3-Fab-bound, 3-“up”, asymmetric structure; **C)** the 3-Fab-bound, 2-“up”-1-“down” structure; “A”, “B” and “C” label the three Fab-RBD units where “C” is in the “down” conformation, “A” stacks over “C” and “B” is the Fab-RBD unit pushed away from the 3-fold rotation axis; **D)** the 2-Fab-bound, 2-“up” structure; **E)** A top view of the “B” Fab-RBD unit (Fab, orange; RBD, green) shown in **C)**. **F)** Comparison of the 15033-7 Fab-RBD unit found in the C3 symmetric cryo-EM structure (Fab, blue; RBD, green) with that found in the crystal structure (Fab and RBD, gray). A change in the Fab elbow angle is observed. **G)** The Fab-RBD unit in the C3 symmetric cryo-EM map, showing two views. **H)** Local-refinement map of the 15033-7 Fab-RBD unit, showing two views. In **G)** and **H)** the Fab heavy chain, Fab light chain and the RBD are colored blue, magenta and green, respectively.

**Figure S3. Thermostability of IgGs and tetravalent nAbs.** Thermal melt curves obtained by monitoring the fluorescence of Sypro orange in the presence of 1 μM antibody from 44 - 100 °C and the relative fluorescence intensity baseline shifted to zero. Vertical bars mark the point of inflection in each curve indicating the T_M_2.

**Figure S4. Antibody-mediated neutralization of clinically isolated SARS-CoV-2** Infection of VeroE6 cells by a clinically-isolated SARS CoV2 virus (strain 2019 n-CoV/USA_WA1/2020) was measured over a range [IgG] versus an IgG isotype control antibody using a focal reduction neutralization assay. Relative infection determined from the number of detectable foci was plotted versus log-transformed [IgG] and plots fit to determine IC_50_ values.

**Figure S5. Vulnerabilities of antibody-mediated neutralization to potential escape mutants. (A)** RBD residues individually mutated to alanine are indicated by spheres and labeled with the residue number. Those variants that exhibited reduced infection >75% relative to the wt are shown in dark grey and were not included in the analysis; those that exhibited at least 25% infectivity but could be neutralized >95% by 50 nM IgG 15033 are shown in light grey and those that retained infectivity but exhibited <95% neutralization of infection by 50 nM IgG 15033 are shown in red. **(B)** Residue Phe486 and Fab CDR residues within 4 Å of it are shown from the crystal structure of the complex of Fab 15033-7 and the SARS CoV-2 RBD. **(C)** Surface view of Fab 15033-7 in complex with the RBD reveals that residues in **(B)** form a hydrophobic pocket between the heavy and light chain, into which Phe486 inserts.

**Figure S6. A negative stain micrograph of the Fab-IgG-S-protein complex** The Fab-IgG-S-protein complexes are indicated by open brackets. Unbound Fab-IgG molecules are indicated by white arrowheads. The micrograph is contrast-inverted to help visualization.

**Figure S7. Negative stain EM particle images of the Fab-IgG-S-protein complexes** Each panel contains a Fab-IgG-S-protein complex observed in negative stain EM micrographs. Each particle is aligned so that the spike portion is upright. The images are contrast-inverted to help visualization.

