## Supplemental Information for "Tetravalent SARS-CoV-2 Neutralizing Antibodies Show Enhanced Potency and Resistance to Escape Mutations"

**Table S1. X-ray Data collection and refinement statistics**

| Structure | RBD-Fab 15033 | RBD-Fab 15033-7 |
|---:|---:|---:|
| **PDB** | **7KLG** | **7KLH** |
| **Data collection** | | |
| Wavelength (Å) | 1.522 | 1.522 |
| Resolution range (Å) | 47.99 - 3.2 (3.314 - 3.2) | 49.32 - 3.0 (3.107 - 3.0) |
| Space group | P 63 2 2 | P 63 2 2 |
| Unit cell | 197.1 197.1 211.51 90 90 120 | 197.27 197.27 211.09 90 90 120 |
| Total reflections | | 637956 (62437) |
| Unique reflections | 40507 (3971) | 48942 (4825) |
| Multiplicity | 12.9 (13.3) | 13.0 (12.9) |
| Completeness (%) | 99.42 (99.90) | 99.60 (99.83) |
| Mean I/sigma(I) | 8.78 (1.11) | 13.15 (0.93) |
| Wilson B-factor | 92.69 | 95.02 |
| R-merge | 0.2504 (2.626) | 0.1579 (2.764) |
| R-meas | 0.2606 (2.726) | 0.1644 (2.88) |
| R-pim | 0.07087 (0.7218) | 0.04501 (0.7978) |
| CC1/2 | 0.998 (0.585) | 0.999 (0.42) |
| CC* | 0.999 (0.859) | 1 (0.769) |
| **Refinement Statistics** | | |
| Reflections used in refinement | 40303 (3969) | 48784 (4821) |
| Reflections used for R-free | 2018 (198) | 2440 (241) |
| R-work | 0.2754 (0.4048) | 0.2599 (0.4154) |
| R-free | 0.2960 (0.4220) | 0.2881 (0.4343) |
| CC(work) | 0.925 (0.647) | 0.947 (0.544) |
| CC(free) | 0.915 (0.583) | 0.923 (0.478) |
| Number of non-hydrogen atoms | 9717 | 9705 |
| macromolecules | 9689 | 9677 |
| ligands | 28 | 28 |
| Protein residues | 1270 | 1270 |
| RMS bond length (Å) | 0.004 | 0.003 |
| RMS bond angle (°) | 0.72 | 0.59 |
| Ramachandran favored (%) | 95.15 | 96.34 |
| Ramachandran allowed (%) | 4.69 | 3.58 |
| Ramachandran outliers (%) | 0.16 | 0.08 |
| Rotamer outliers (%) | 0 | 0 |
| Clashscore | 4.25 | 2.83 |
| Average B-factor | 96.78 | 103.02 |
| macromolecules | 96.75 | 102.97 |
| ligands | 108.19 | 120.94 |

**Table S2. Cryo-EM data collection and image processing**

| **Structure** | spike, 2x Fab 15033-7 2-"up"-1-"down" | spike, 3x Fab 15033-7 3-"up", C3-symmetric | spike, 3x Fab 15033-7 3-"up", asymmetric | spike, 3x Fab 15033-7 2-"up", 1-"down" |
|---|---|---|---|---|
| **EMDB ID** | **EMD-22925** | **EMD-22926** | **EMD-23064** | **EMD-23065** |
| **PDB ID** | **7KMK** | **7KML** | **7KXJ** | **7KXK** |
| **Data collection** | | | | |
| Electron microscope | Titan Krios G3 | | | |
| Camera | Falcon 4EC | | | |
| Voltage | 300 kV | | | |
| Nominal magnification | 75000x | | | |
| Calibrated physical pixel size (Å) | 1.03 | | | |
| Total exposure (e/Å2) | 38 | | | |
| Number of fractions | 30 | | | |
| Movies collected | 6431 | | | |
| **Image Processing and map refinement** | | | | |
| Motion correction software | cryoSPARC v2 | | | |
| CTF estimation software | Gctf | | | |
| Particle selection software | cryoSPARC v2 | | | |
| Classification and refinement software | cryoSPARC v2 | | | |
| symmetry | C1 | C3 | C1 | C1 |
| Global resolution (Å) | 4.2 | 3.8 | 6.4 | 5.0 |
| Particles used in final reconstruction | 88121 | 95627 | 19743 | 32257 |
| **Structural refinement** | | | | |
| Modeling software | Coot, Rosetta, Phenix | | | |
| RMS bond length (Å) | 0.019 | 0.018 | 0.020 | 0.018 |
| RMS bond angle (̊) | 1.84 | 1.88 | 1.91 | 1.84 |
| MolProbity score | 0.77 | 0.68 | 0.75 | 0.77 |
| Clash score | 0.79 | 0.53 | 0.79 | 0.88 |
| Ramachandran favored | 97.95% | 98.14% | 98.38% | 98.03% |
| Ramachandran outliers | 0.08% | 0.00% | 0.07% | 0.07% |
| Rotamer outliers | 0.17% | 0.39% | 0.18% | 0.42% |
| Cbeta outliers | 0.03% | 0.00% | 0.05% | 0.07% |
| Average B factor (protein) | 352 | 259 | 649 | 505 |
| Average B factor (glycan) | 381 | 248 | 660 | 412 |

### Figure S1

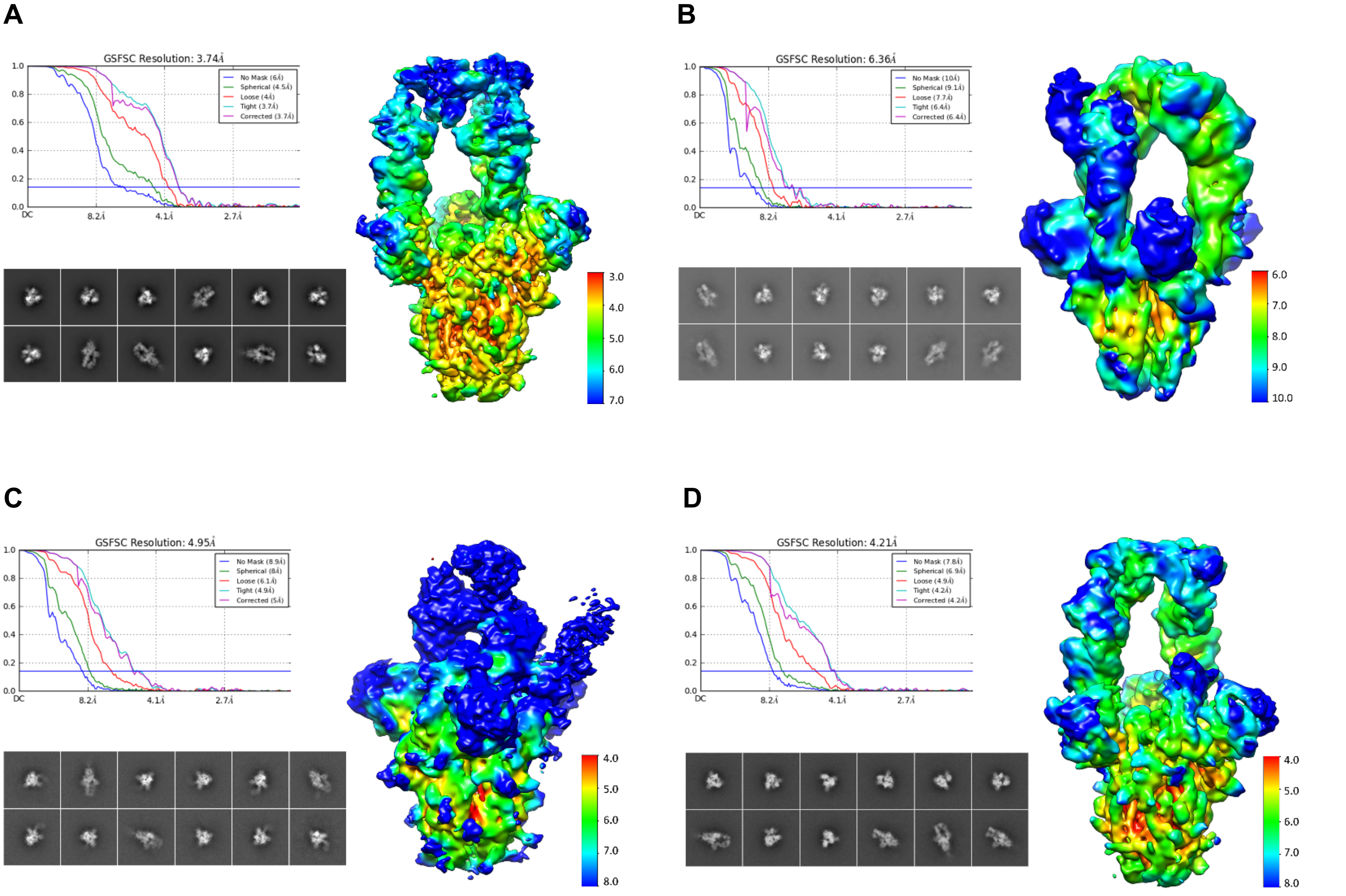


**Figure S1. 2D class averages and resolution plots of the four 15033-7 Fab-S-protein cryo-EM structures** The GSFSC curve, selected 2D class averages and the local resolution map are shown for each of the four structures: **A)** the 3-Fab-bound, 3-"up", C3 symmetric structure; **B)** the 3-Fab-bound, 3-"up", asymmetric structure; **C)** the 3-Fab-bound, 2-"up"-1-"down" structure; **D)** the 2-Fab-bound, 2-"up" structure.

**Figure S2. Cryo-EM maps showing only Fabs and RBDs in the density** (on next page) **A-D)** The 15033-7 Fabs (colored ribbons) are shown in the cryo-EM maps for each of the four structures: **A)** the 3-Fab-bound, 3-"up", C3 symmetric structure; **B)** the 3-Fab-bound, 3-"up", asymmetric structure; **C)** the 3-Fab-bound, 2-"up"-1-"down" structure; "A", "B" and "C" label the three Fab-RBD units where "C" is in the "down" conformation, "A" stacks over "C" and "B" is the Fab-RBD unit pushed away from the 3-fold rotation axis; **D)** the 2-Fab-bound, 2-"up" structure; **E)** A top view of the "B" Fab-RBD unit (Fab, orange; RBD, green) shown in **C)**. **F)** Comparison of the 15033-7 Fab-RBD unit found in the C3 symmetric cryo-EM structure (Fab, blue; RBD, green) with that found in the crystal structure (Fab and RBD, gray). A change in the Fab elbow angle is observed. **G)** The Fab-RBD unit in the C3 symmetric cryo-EM map, showing two views. **H)** Local-refinement map of the 15033-7 Fab-RBD unit, showing two views. In **G)** and **H)** the Fab heavy chain, Fab light chain and the RBD are colored blue, magenta and green, respectively.

**Figure S2**

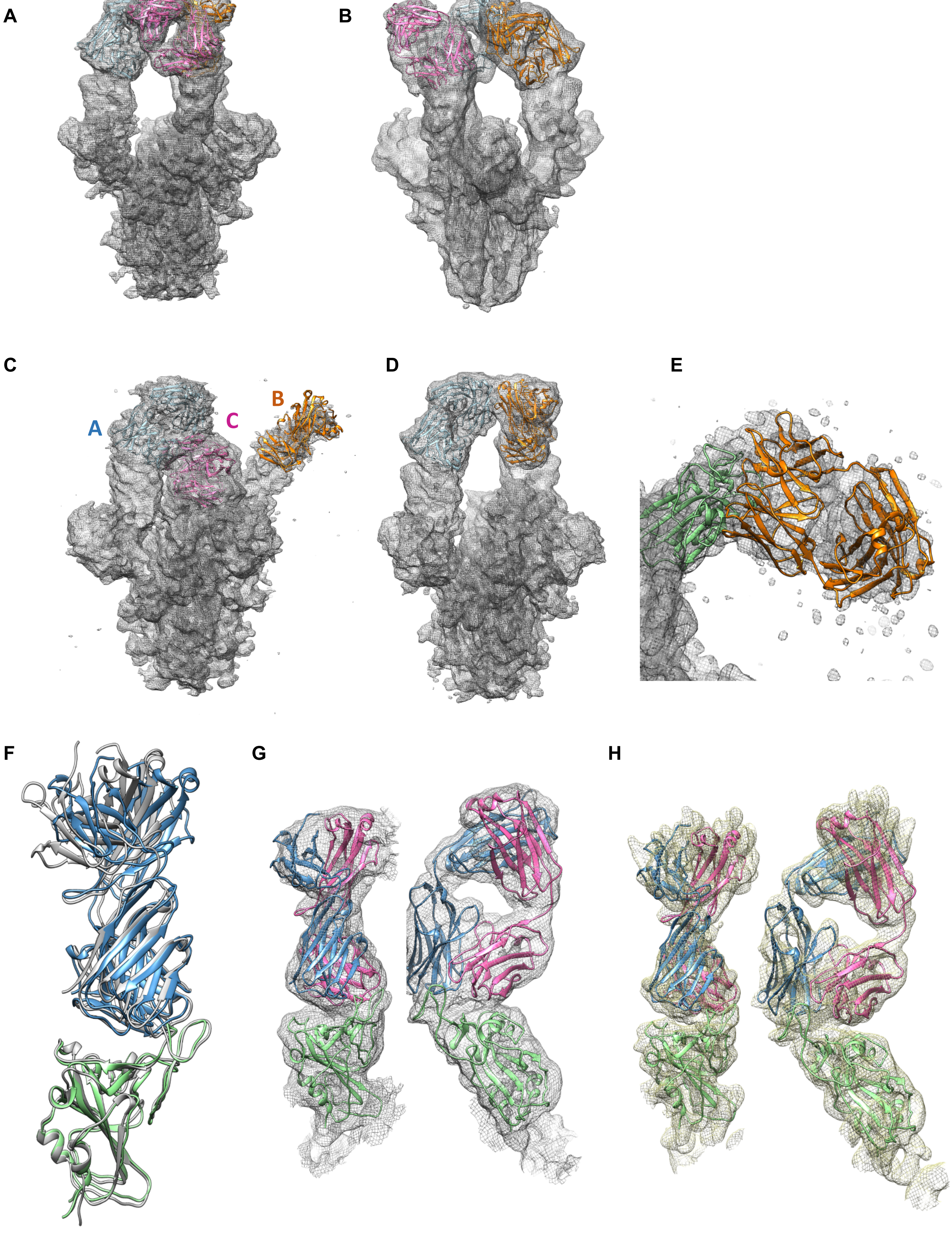

**Figure S3**

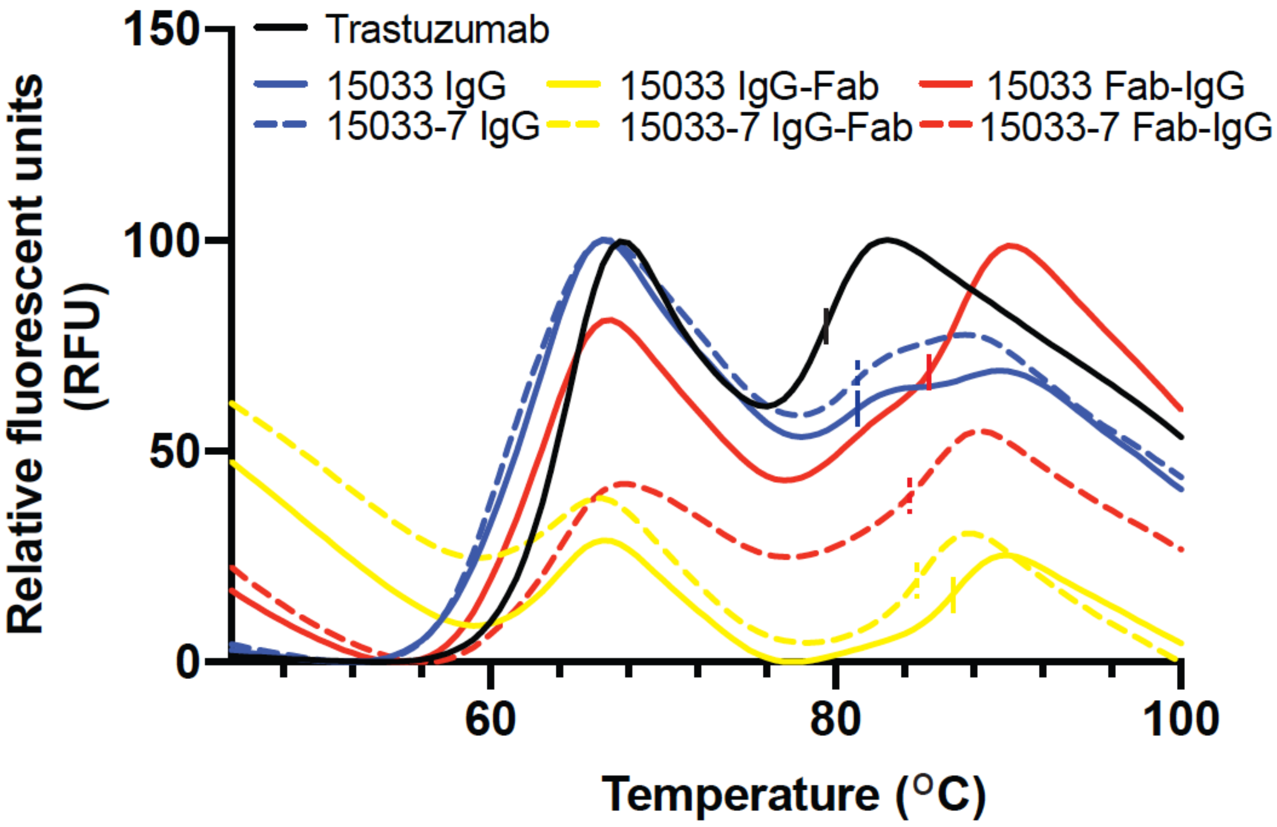


**Figure S3. Thermostability of 15033 IgG and tetravalent antibodies and matured variants** Thermal melt curves obtained by monitoring the fluorescence of Sypro orange in the presence of 1 μM antibody from 44 - 100 °C and the relative fluorescence intensity baseline shifted to zero. Vertical bars mark the point of inflection in each curve indicating the $T_M2$.

.

### Figure S4

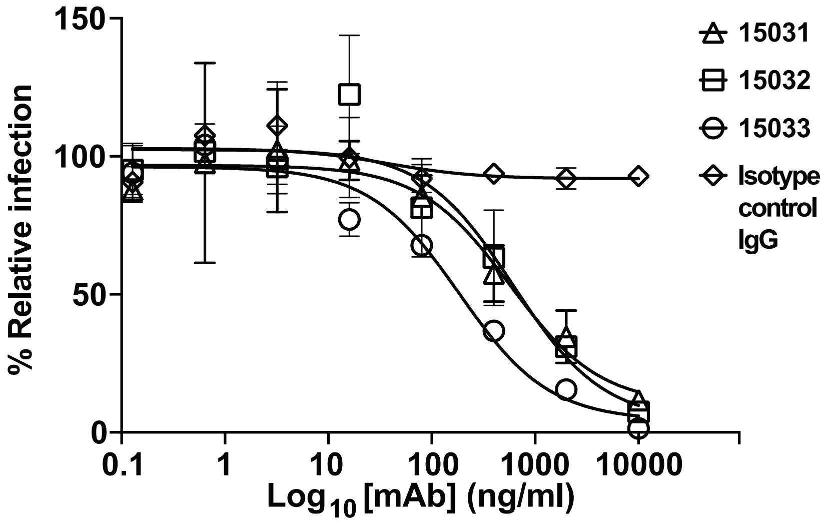


**Figure S4. Antibody-mediated neutralization of clinically isolated SARS CoV2 virus** Infection of VeroE6 cells by a clinically-isolated SARS CoV2 virus (strain 2019 n-CoV/USA_WA1/2020) was measured over a range [IgG] versus an IgG isotype control antibody using a focal reduction neutralization assay. Relative infection determined from the number of detectable foci was plotted versus log-transformed [IgG] and plots fit to determine $IC_{50}$ values.

**Figure S5**

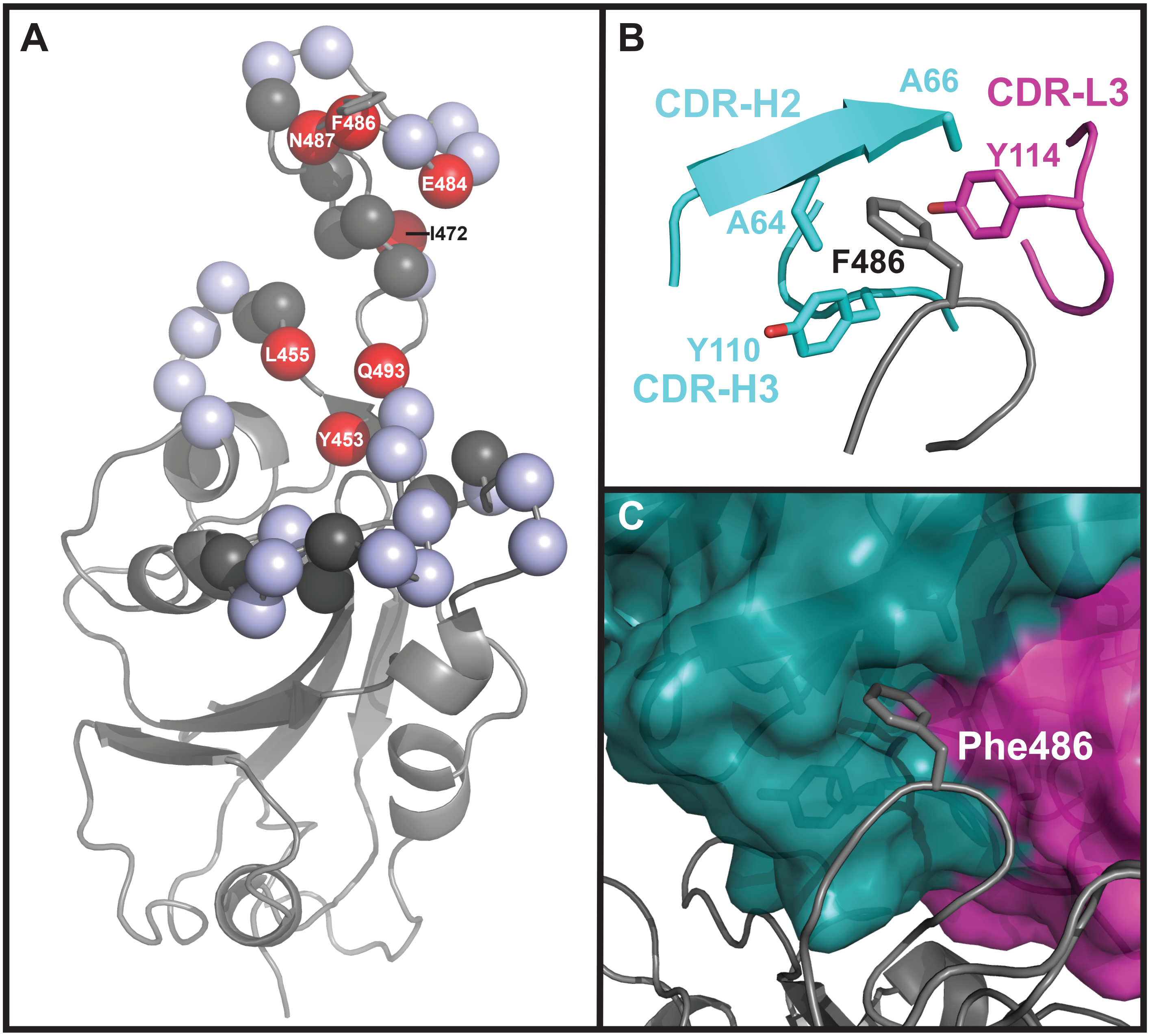


**Figure S5. Vulnerabilities of antibody-mediated neutralization to potential escape mutants. (A)** RBD residues individually mutated to alanine are indicated by spheres and labeled with the residue number. Those variants that exhibited reduced infection >75% relative to the wt are shown in dark grey and were not included in the analysis; those that exhibited at least 25% infectivity but could be neutralized >95% by 50 nM IgG 15033 are shown in light grey and those that retained infectivity but exhibited <95% neutralization of infection by 50 nM IgG 15033 are shown in red. **(B)** Residue Phe486 and Fab CDR residues within 4 Å of it are shown from the crystal structure of the complex of Fab 15033-7 and the SARS CoV-2 RBD. **(C)** Surface view of Fab 15033-7 in complex with the RBD reveals that residues in **(B)** form a hydrophobic pocket between the heavy and light chain, into which Phe486 inserts.

**Figure S6**

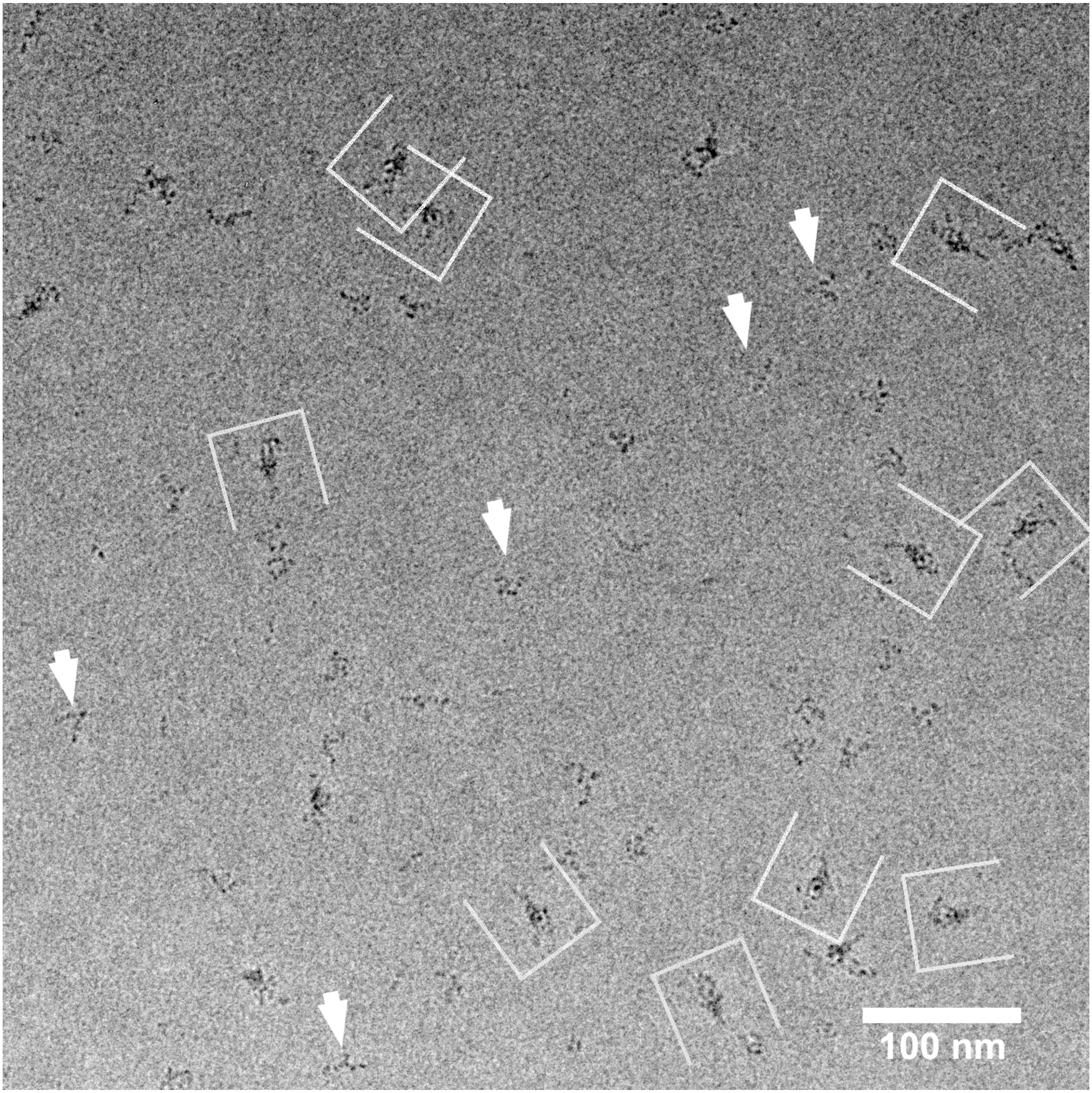


**Figure S6. A negative stain micrograph of the Fab-IgG-S-protein complex** The Fab-IgG-S-protein complexes are indicated by open brackets. Unbound Fab-IgG molecules are indicated by white arrowheads. The micrograph is contrast-inverted to help visualization.

**Figure S7**

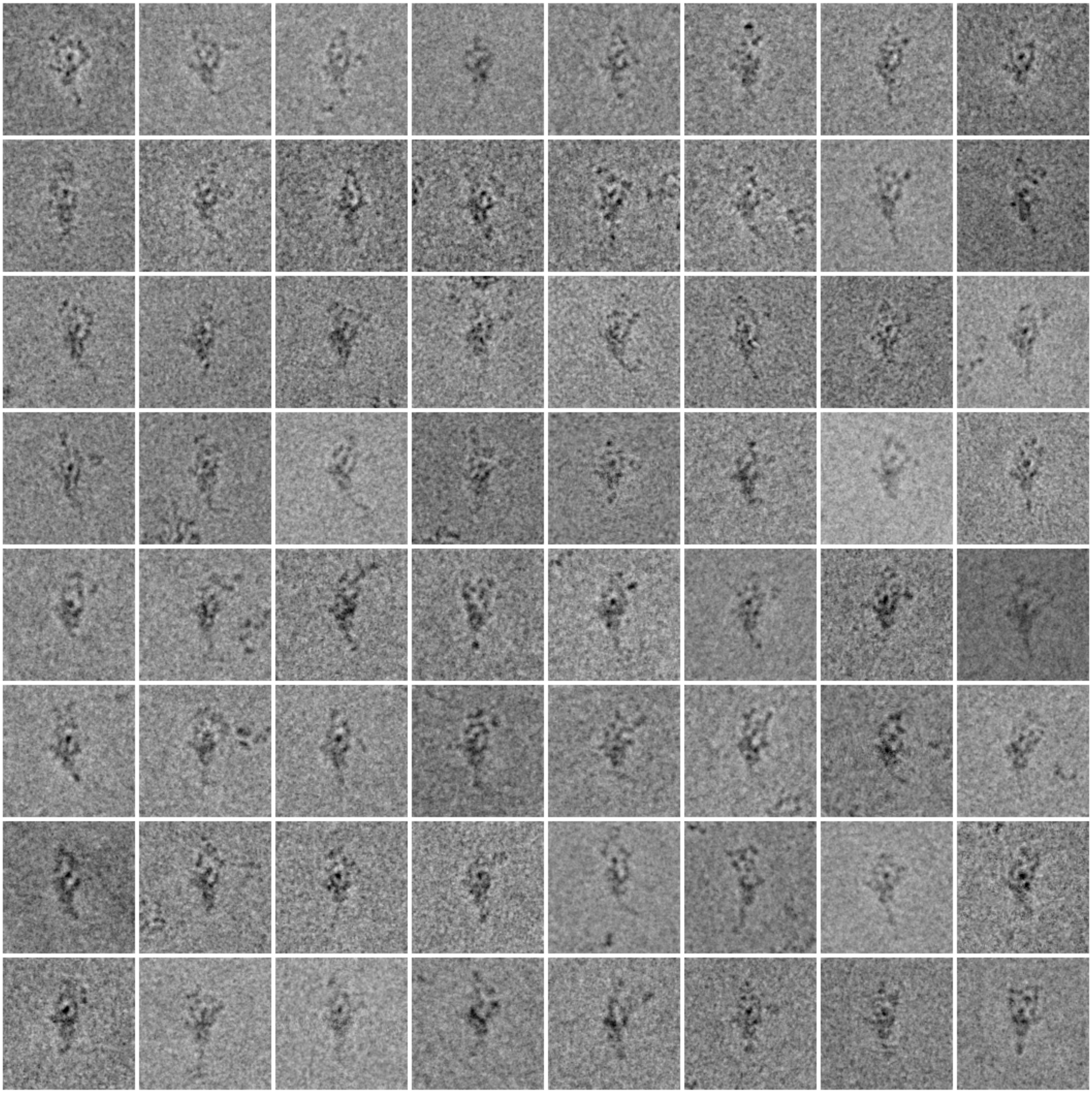

**Figure S7. Negative stain EM particle images of the Fab-IgG-S-protein complexes** Each panel contains a Fab-IgG-S-protein complex observed in negative stain EM micrographs. Each particle is aligned so that the spike portion is upright. The images are contrast-inverted to help visualization.
